# Transfer of disulfide bond formation modules via yeast artificial chromosomes promotes the expression of heterologous proteins in *Kluyveromyces marxianus*

**DOI:** 10.1101/2023.11.30.569359

**Authors:** Pingping Wu, Wenjuan Mo, Tian Tian, Kunfeng Song, Yilin Lyu, Haiyan Ren, Jungang Zhou, Yao Yu, Hong Lu

## Abstract

*Kluyveromyces marxianus* is a food-safe yeast with great potential for producing heterologous proteins. Improving the yield in *K. marxianus* remains a challenge, while incorporating large-scale functional modules poses a technical obstacle in engineering. To address these issues, linear and circular yeast artificial chromosomes of *K. marxianus* (KmYACs) were constructed and loaded with disulfide bond formation modules from *Pichia pastoris* or *K. marxianus*. These modules contained up to 7 genes with a maximum size of 15 kb. KmYACs carried telomeres either from *K. marxianus* or *Tetrahymena*. KmYACs were transferred successfully into *K. marxianus* and stably propagated without affecting the normal growth of the host, regardless of the type of telomeres and configurations of KmYACs. KmYACs increased the overall expressions of disulfide bond formation genes and significantly enhanced the yield of various heterologous proteins. In high-density fermentation, the use of KmYACs resulted in a glucoamylase yield of 16.8 g/L, the highest reported level to date in *K. marxianus*. Transcriptomic and metabolomic analysis of cells containing KmYACs suggested increased FAD biosynthesis, enhanced flux entering the TCA cycle and a preferred demand for lysine and arginine as features of cells overexpressing heterologous proteins. Consistently, supplementing lysine or arginine further improved the yield. Therefore, KmYAC provides a powerful platform for manipulating large modules with enormous potential for industrial applications and fundamental research. Transferring the disulfide bond formation module via YACs proves to be an efficient strategy for improving the yield of heterologous proteins, and this strategy may be applied to optimize other microbial cell factories.

**Impact Statement:** In this study, yeast artificial chromosomes of *K. marxianus* (KmYACs) were constructed and successfully incorporating modules for large-scale disulfide bond formation. KmYACs were stably propagated in *K. marxianus* without compromising the normal growth of the host, irrespective of the selection of telomeres (either *Tetrahymena* or *K. marxianus*) and configuration (either linear or circular). KmYACs notably enhanced the expressions of various heterologous proteins, with further yield improvement by supplementing lysine or arginine in the medium. Our findings affirm KmYAC as a robust and versatile platform for transferring large-scale function modules, demonstrating immense potential for both industrial applications and fundamental research.

## Introduction

*Kluyveromyces marxianus* is a budding yeast belonging to the *Saccharomyces* subclade within hemiascomycetes. Due to its long-standing safe association with human food, particularly dairy products, *K. marxianus* has been granted GRAS (Generally Regarded As Safe) and QPS (Qualified Presumption of Safety) status in the United States and Europe, respectively ^1^. It has been approved as a new food raw material and a new feed additive in China in 2023. Therefore, *K. marxianus* is a native food-safe yeast and is an ideal host for producing food proteins. Besides food-grade safety of *K. marxianus*, this species has many excellent properties that are suitable for industrial production, including thermotolerance^2^, high biomass, high growth rate^1^ and broad substrate spectrum^3^. Starting from the 1980s, *K. marxianus* emerges as an excellent microbial cell factory for producing diverse recombinant proteins and chemicals, such as glycosidase^4^, lignocellulolytic enzymes^5^, virus-like particles^6^, bioethanol^7^, xylitol^8^, aromatic amino acid^9^ and flavour metabolites^10^.

With the expansion of *K. marxianus* application scenarios, a series of genetic tools for the engineering of *K. marxianus* were developed, including episomal plasmids^11–12^, multiple-cassette integration^13^, CRISPR/Cas9 system^14^, Cre-loxP system^15^, flux balance analysis (FBA) model^16^, and genome-scale metabolic model^17^. Despite these achievements, the incorporations of large-scale functional modules remains an obstacle. Large-scale modules can be integrated into the genome through homologous recombination, but the efficiency of integration is low. So far, the largest size for one-piece fragment integration was 5 kb^18^, and that for overlapping fragments integration was below 15 kb^13^. In addition, the expression of the function module is highly dependent on the integrated locus and may interfere with the expressions of native genes around the locus^19–20^. Episomal plasmids, either multicopy or centromeric, can be applied in transferring large functional modules. The most commonly used multicopy plasmids in *K. marxianus* were derived from the endogenous pKD1 plasmid of *K. lactis*^5, 11^. However, the copy number of pKD1-derived plasmids is not stable^5^, making them only suitable for overexpression rather than the stable expression of genes in the functional module. The capacity of centromeric plasmids is limited. Meanwhile, the loss rates of centromeric plasmids were approximately 1%-2% per generation^15^, so they are not suitable for long-term culturing without selective pressure. Regarding the problem of integration and episomal plasmids, yeast artificial chromosome (YAC) provides a new option. YACs are shuttle (*E. coli* and *S.cerevisiae*) vectors containing essential functional elements of a typical chromosome, including autonomously replicating sequences (ARS), centromere (CEN) and telomere (TEL)^21^. YACs are characterized by their extremely large capacity to accept exogenous DNA (up to 3 Mb)^22^. After exceeding 50 kb in size, YACs exhibit high stability comparable to natural chromosomes, with a loss rate of ∼0.3% per generation, which is superior to centromeric plasmids^21,23^. YACs are independent of natural chromosomes, avoiding mutual interference between native genes and loaded modules caused by integration. After its invention in 1983, YACs were originally used for the cloning and manipulation of large DNA segments from higher organisms, especially those from mammalians^24–25^. In recent years, YACs received attention from the field of yeast synthetic biology and have been successfully applied as a vector to transfer large functional modules into *S. cerevisiae* and other unconventional yeast. For example, YAC was applied to transfer cellobiose phosphorolysis and xylose consumption modules (∼12.2 kb) into *Yarrowia lipolytic*a^26^. In *S. cerevisiae*, the entire flavonoid pathway was cloned in YACs to generate a small library for screening flavonoid intermediates or derivatives^27^. YAC containing xylose utilization enzymes that enhance xylose utilization was constructed in *S. cerevisiae*^28^. However, no YAC has been constructed for *K. marxianus* so far.

Currently, more than 50 heterologous proteins have been successfully expressed in *K. marxianus*. The highest yield of a heterologous protein in *K. marxianus* was 12.2g/L^29^, which is higher than that of *S. cerevisiae* (5 g/L)^30^, but is significantly lagging behind that of *Pichia pastoris* (22 g/L)^31^. This indicates that there is still significant room to improve the yield of *K. marxianus*. One crucial step in the folding and maturation of proteins that enter the secretory system is the formation of native disulfide bonds between two cysteine residues. The process involves the oxidation of substrate proteins by Protein disulfide isomerase (Pdi1), which in turn is oxidized by endoplasmic reticulum oxidase (Ero1 and Erv2) ^32^. Transferring disulfide bond formation module genes is an efficient strategy to increase the production of heterologous protein. Overexpression of native *PDI1* at the *GOS1* locus showed a 24% increase in α-amylase production in *S. cerevisiae*^33^. Duplication of either *KlERO1* or *KlPDI1* resulted in an approximately 15-fold higher yield of Human Serum Albumin secreted by *K. lactis*^34^. Simultaneous co-expression of native *ERO1* and *PDI1* resulted in approximately 30% higher enzyme yields of lipase r27RCL in *P. pastoris*^35^. The effect of transferring genes involved in disulfide bond formation modules on protein production in *K. marxianus* has not been evaluated yet. It remains unclear whether transferring additional genes from the disulfide bond formation modules, beyond the few genes mentioned above, would further enhance the protein yield. Furthermore, the compatibility of the introduced genes with the native genes is also unknown.

In this study, yeast artificial chromosomes for *K. marxianus* (KmYACs) were constructed and used to transfer disulfide bond formation modules into the organism. The module contained at most 7 genes, with a size of 15 kb. KmYAC was stably inherited in *K. marxianus* as a single copy. The *P. pastoris* genes introduced by KmYAC were actively expressed in vivo and were compatible with the native genes of disulfide bond isomerization pathway in *K. marxianus*.Transfer of the disulfide bond formation module significantly enhances the secretory expressions of various heterologous proteins in *K. marxianus*, with the highest level reaching 16.8 g/L, the highest level reported to date in *K. marxianus*.

## Results

### Design and construction of KmYACs carrying disulfide bond formation modules

The KmYAC shuttle vector contained a replication origin *ARS1*, a centromere *CEN5*, two inverted telomeres (either from *Tetrahymena* or *K. marxianus*), and three selection marker *HIS3*, *TRP1* and *HphMX4* (Figure 1). To construct a linear KmYAC, the shuttle vector was digested with *Bam*H I and *Not* I. The purpose of *Bam*H I digestion was to release the left and right arms of KmYAC. Exogenous fragments (two cassettes were used in this study) were ligated to arms at the *Not* I site using Gibson assembly in vitro. The *Not* I site was in frame with the ORF of *HphMX4*, and the insertion would abolish the function of *HphMX4*. The ligation product was transformed into *K. marxianus* cells and transformants containing the linear KmYAC were selected. The same procedure was followed to construct circular KmYAC, except that the *Bam*H I digestion was omitted to maintain the circular formation of KmYAC (Figure 1).

**Figure 1.**
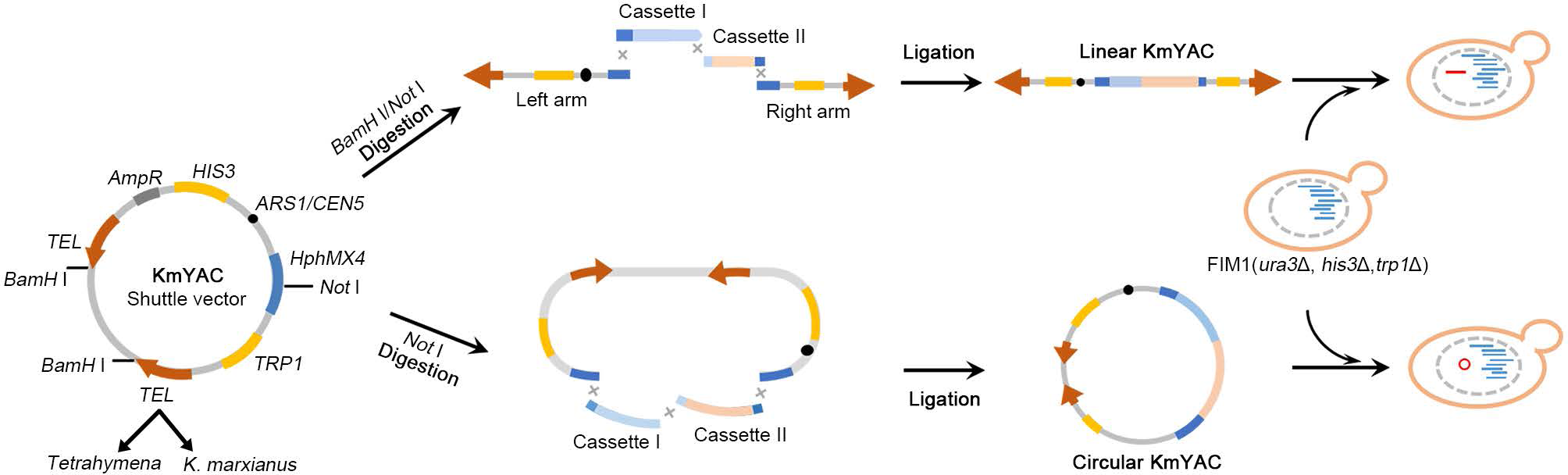
Schematic map of KmYACs construction. The KmYAC shuffle vector was composed of a replication origin (*ARS1*), a centromere (*CEN5*), two inverted telomeres (*TEL*) from either *Tetrahymena* or *K. marxianus*, and three selection markers (*HIS3*, *TRP1* and *HphMX4*). KmYAC shuffle vectors were digested and ligated with exogenous fragments by Gibson assembly in vitro to construct linear or circular KmYAC. The ligation product was then transformed into *K. marxianus* cells and transformants containing KmYACs were selected.

To improve the yield of heterologous proteins in *K. marxianus*, we designed our module to include genes encoding essential enzymes implicated in disulfide bond formation, including Ero1, Erv2, Pdi1 and Mpd1 (Figure 2A). Ero1 and Erv2 are ER oxidases that catalyze de novo disulfide bond formation and transfer bonds to Pdi1, oxidized Pdi1 then introduces disulfide bonds into reduced substrates^36–37^. Mpd1 is a member of the Pdi1 family and can fully compensate for the deletion of Pdi^38–39^. Downstream of the disulfide bond formation, Ypt1 is a GTPase required for vesicle docking and vesicle targeting during ER to Golgi trafficking. Given its importance in protein secretion, *YPT1* was also included in our designed module (Figure 2A). A total of three modules, namely P5, P7, and K5 were designed (Figure 2B). Module P5 and P7 comprised five and seven genes from *P. pastoris*, including three *MPD1* homologues named *PpMPD1-1*, *PpMPD1-2*, and *PpMPD1-3*. Module K5 included five genes from *K. marxianus*. P5, P7, and K5 modules were loaded into linear KmYAC containing either *K. marxianus* telomeres or *Tetrahymena* telomeres, resulting in six assemblies. Three modules were loaded into circular KmYAC containing *K. marxianus* telomeres, resulting in three assemblies (Figure 2C). The average fidelity of linear KmYACs (23.9%) was higher than that of circular KmYACs (6.9%), suggesting linear configuration is more suitable for assembly. The average assembly fidelity of linear KmYACs containing *K. marxianus* telomeres (41.6%) was higher than that of those containing *Tetrahymena* telomeres (6.1%), suggesting the native *K. marxianus* telomeres promote accurate assembly (Figure 2C).

**Figure 2.**
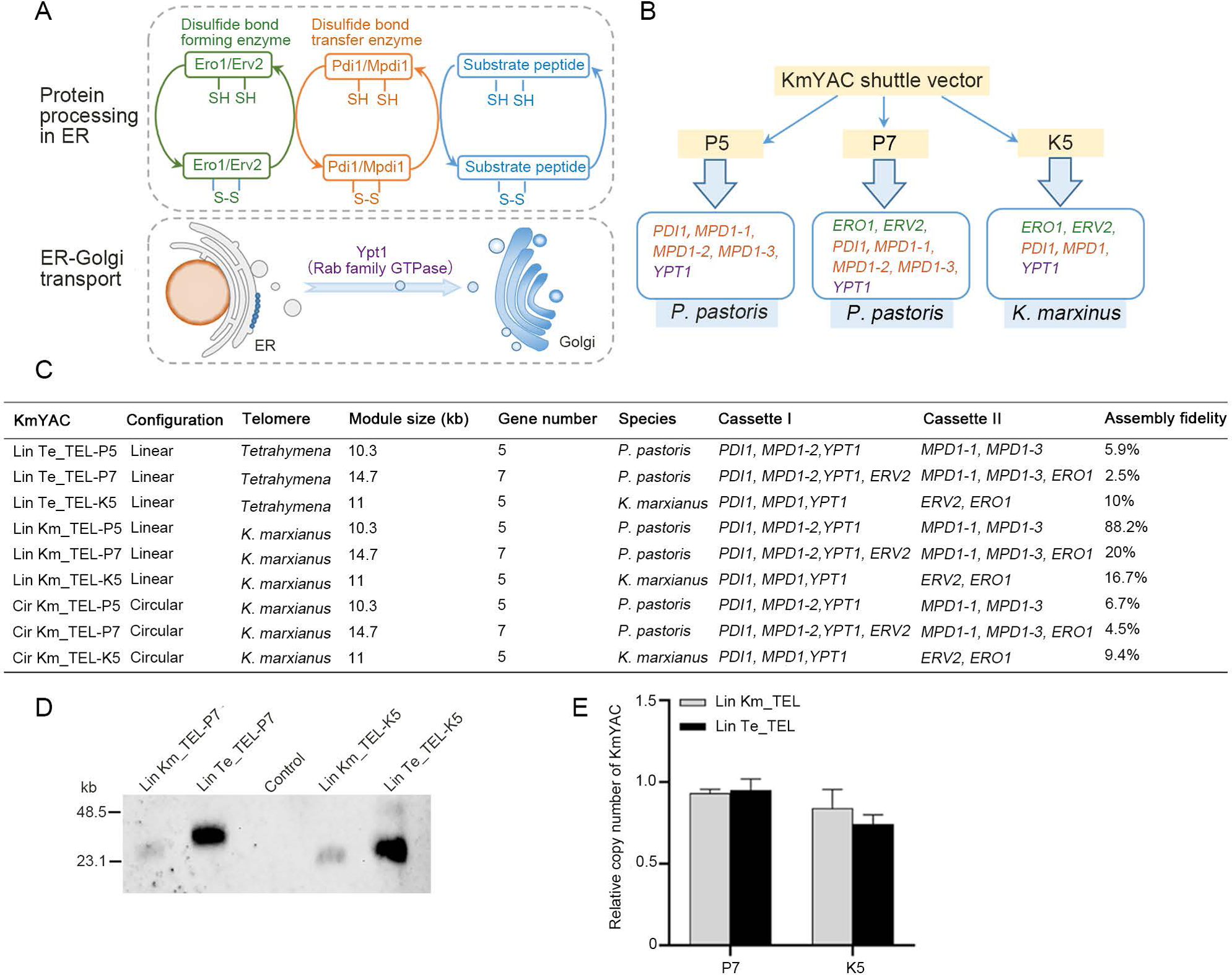
The construction of KmYACs carrying disulfide bond formation modules. (A) Genes in the disulfide bond formation modules. Genes encoding ER oxidase (Ero1, Erv2), protein disulfide isomerase (Pdi1, Mpd1), and Rab family GTPase (Ypt1) were included in the module. (B) Composition of P5, P7, and K5 modules. Each module contained genes either from *P. pastoris* or *K. marxianus*. Module was loaded onto KmYAC shuttle vector to construct KmYAC. (C) The characteristics of nine KmYACs. Assembly fidelity was calculated as the ratio of positive clones containing expected KmYACs to total clones. (D) Southern blot of KmYACs. (E) The copy number of KmYACs per cell. Copy number of KmYAC were calculated relative to that of endogenous *LEU2*. The values presented represent the mean ± S.D. (n=3).

Southern blot was performed to confirm the presence of KmYACs in *K. marxianus*. The probe detected bands corresponding to both linear and circular KmYACs (Figure 2D, Figure S1). Notably, the size of the linear KmYACs was larger than the predicted size, possibly due to telomere extension. Moreover, the linear KmYAC that contained *Tetrahymena* telomeres was larger than the one containing *K. marxianus* telomeres, suggesting additional telomeric sequences were added at the heterologous telomeres (Figure 2D). The difference in intensity between the bands of linear and circular KmYACs might be attributed to differences in loading rather than variations in copy numbers per cell. This was supported by the qPCR results which indicated that the copy numbers of both linear and circular KmYACs per cell were close to 1 (Figure 2E).

### Stabilities of KmYACs and their effects on growth

The mitotic stabilities of linear and circular KmYACs were investigated by culturing cells without selective pressure (Figure 3A). At 30℃, the average loss rate of KmYACs was ∼0.48% per generation, which was comparable to the loss rate of YACs in *S. cerevisiae* (∼0.3%) and was significantly lower than that of YACs in *Y. lipolytica* (∼6%)^21, 23, 26^. At 40℃, Lin Km-TEL-K5 exhibited a relatively high loss rate (7.7% per generation), while other KmYACs still showed high stability with an average loss rate of 0.75% per generation. The high stability of most KmYACs at elevated temperatures might be associated with the thermotolerance characteristics of *K. marxianus*^40^.There was no statistical difference between the stabilities of KmYACs carrying *K. marxianus* telomeres and those carrying *Tetrahymena* telomeres, suggesting heterologous telomeres support faithful segregation as well as native telomeres. Additionally, the configurations of KmYACs had no significant effect on stability, as circular KmYACs and linear KmYACs shared similar stabilities. The high stabilities of KmYACs provide advantages for long-term fermentation in various temperatures without selective pressure.

**Figure 3.**
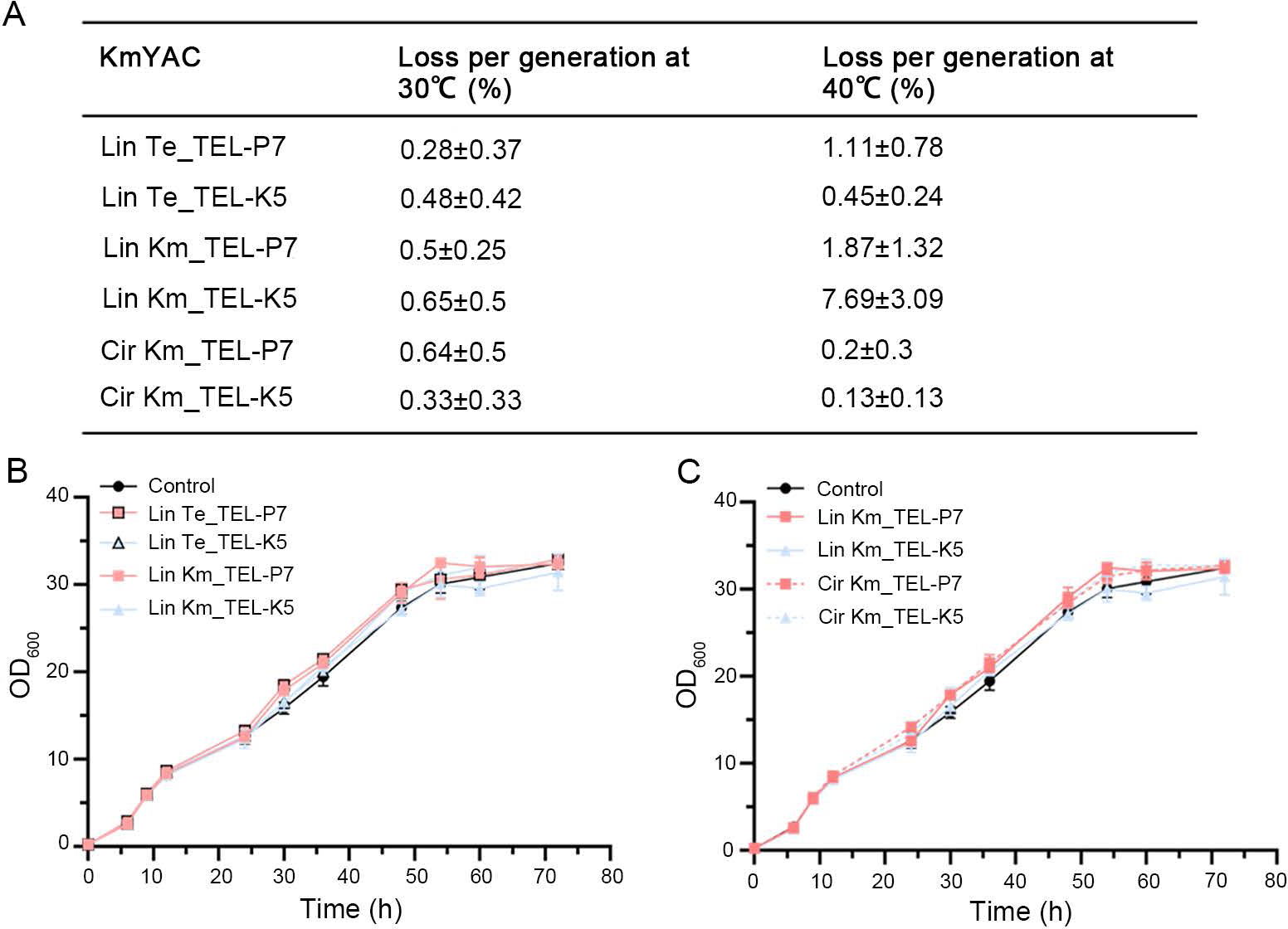
Stabilities of KmYACs and their effects on cell growth. (A) Loss percentage of the KmYAC per generation in cells grown without selective pressure at 30℃ or 40 ℃. Values represent mean ± S.D. from three parallel cultures. (B) Growth curves of cells containing linear KmYACs with telomeres either from *K. marxianus* or *Tetrahymena* at 30℃. Cells without any KmYAC served as a control. Values represent mean ± SD (n=3). (C) Growth curves of cells containing linear or circular KmYACs.

The growth rate is a crucial factor in the production of microbial cell factories. Therefore, the effects of KmYACs on the growth of host cells were monitored. The growth curve of control cells was the same as cells containing linear KmYACs with either *K. marxianus* telomeres or *Tetrahymena* telomeres (Figure 3B). Meanwhile, growth curves of cells containing either linear or circular KmYACs were the same as that of control cells. Therefore, the presence of KmYACs, regardless of their configuration or type of telomere, did not affect the normal growth of host cells (Figure 3C).

### KmYACs carrying disulfide bond formation modules improve expressions of heterologous proteins

Next, the effects of KmYACs on the secretory expressions of heterologous proteins were investigated. A multicopy plasmid expressing glucoamylase BadGLA was transformed into cells with KmYACs carrying diverse disulfide bond formation modules. Compared to the control cells, the secretory activities of BadGLA were significantly increased in cells containing KmYACs (Figure 4A). The magnitude of increase ranged from 23% to 129%, with the highest increase observed in cells containing the linear KmYAC with *K. marxianus* telomeres and P5 module (Lin Km_TEL-P5). Three linear KmYACs were chosen to evaluate their effects on the expressions of two other enzymes. Compared to the control, the linear KmYAC carrying the P5, P7, and K5 modules increased the activities of glucoamylase TeGlaA by 87%, 65%, and 123%, respectively (Figure 4B). The same set of KmYACs increased the activities of the β-1,4-endoxynlanase Xyn-CDBFV by 90%, 92%, and 56%, respectively (Figure 4C). The improved yield of TeGlaA and Xyn-CDBFV was confirmed by SDS-PAGE analysis (Figure 4D-E). Therefore, the KmYACs carrying disulfide bond formation modules exhibited universality in promoting the expression of heterologous proteins.

**Figure 4.**
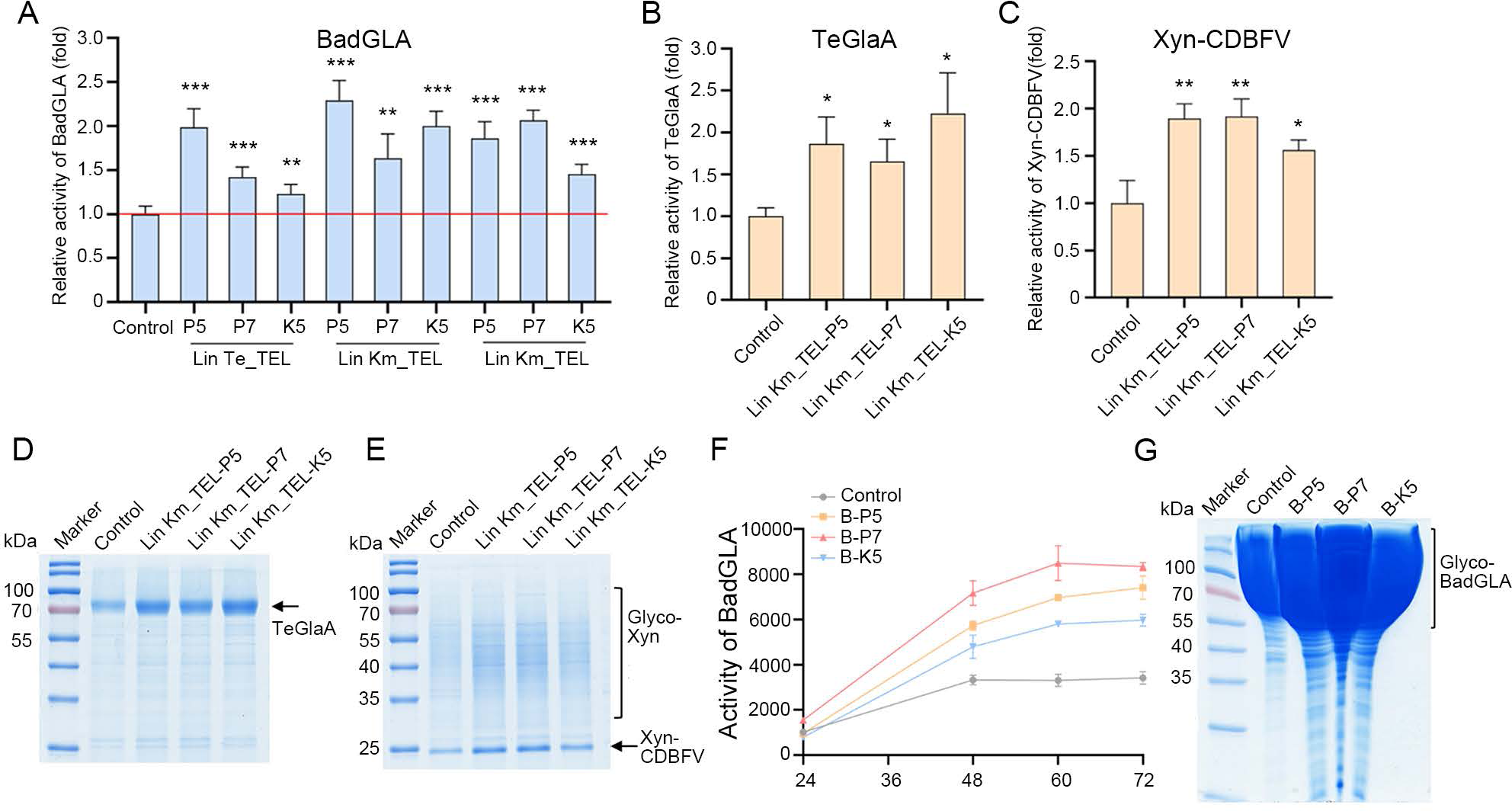
KmYACs promote secretory expression of heterologous proteins. (A-C) The activity of BadGLA (A), TeGlaA (B) or Xyn-CDBFV (C) in the supernant of culture grown with cells containing KmYACs. The cells containing KmYACs were transformed with a plasmid expressing the indicated heterologous protein, while cells without KmYACs were transformed with the plasmid as a control. Transformants were cultivated for 72 hours and the activity in the supernatant was measured. The activity from control cells was designated as unit 1. The values represent the mean ± S.D. (n=3). * *p* <0.05, ** *p* <0.01, \*\*\**p* <0.001. (D-E) SDS-PAGE of the superanatant described in (B) and (C). (F) Activity curves of BadGLA in supernatant of culture grown in a 5L fermentor. Values represent the mean from three technical repeats. (G) SDS-PAGE of 16 μL supernatant of culture grown in 5L fermentor for 72 h. SDS-PAGE of the samples diluted for 35 times shown in Figure S2. A smear at high molecular weight was proposed to be the hyperglycosylated form of BadGLA, as the smear disappeared after deglycosylation (Figure S3).

We subsequently investigated whether KmYACs promote the yield of heterologous protein in high-density fermentation. For this purpose, we cultivated cells carrying the BadGLA gene and either Lin Km_TEL-P5 (referred to as B-P5), Lin Km_TEL-P7 (referred to as B-P7), or Lin Km_TEL-K5 (referred to as B-K5) in a fed-batch 5L fermentor. After 72 hours of cultivation, the secretory activity of BadGLA in B-P5, B-P7 and B-K5 reached 7414.6 U/mL, 8345.7 U/mL, and 5973.2 U/mL, representing increases of 116.5%, 143.7%, and 74.5% when compared to control cells (Figure 4F). The high-yield production of BadGLA was confirmed by SDS-PAGE analysis (Figure 4G, Figure S2). Based on the specific activity of BadGLA (498.02 U/mg) (Figure S3), the secretory expression of BadGLA in B-P5, B-P7 and B-K5 was determined to be 14.9 g/L, 16.8 g/L, and 12 g/L, respectively. It is noteworthy that the yield of 16.8 g/L represents the highest reported yield of a heterologous protein in *K. marxianus* thus far. In high-density fermentation, the yield of B-P7 was superior to that of B-P5, indicating that a more complete disulfide bond isomerization pathway transferred by KmYAC is more favorable in increasing heterologous protein yield. Additionally, the yields of B-P7 and B-P5 were higher than that of B-K5, suggesting that *P. pastoris* genes work more efficiently in protein synthesis and transportation than the native genes of *K. marxianus*. To confirm the association between the improved yield of heterologous proteins and modules transferred by KmYAC, expressions of module genes in KmYAC were evaluated. B-P7 and B-K5 were collected for analysis after cultivating 48 and 72 h. Compared to a native housekeep gene (*KmSWC4*), *PpERO1*, *PpERV2*, *PpPDI1*, *PpMPD1-1*, *PpMPD1-2*, *PpMPD1-3* and *PpYPT1* of the P7 module in B-P7 were all actively transcribed (Figure 5A). The effect of P7 module on the expressions of homologous genes in *K. marxianus* were evaluated. After 48 h, B-P7 did not show any significant difference in the expression levels of *KmERO1*, *KmERV2*, *KmPDI1*, *KmMPD1*, and *KmYPT1* compared to the control cells (Figure 5B). This trend remained unchanged after 72 h, except for a slight decrease in the expression level of *KmYPT1* observed in B-P7 (Figure 5C). The *P. pastoris* genes introduced by KmYAC were actively expressed in vivo and were compatible with the native genes of disulfide bond isomerization pathway in *K. marxianus*. In the case of KmYAC carrying native *K. marxianus* genes, the expression level of *KmERO1*, *KmERV2*, *KmPDI1*, *KmMPD1* and *KmYPT1* were all increased in the B-K5 compared to control cells. The magnitude of increase ranged from 0.24 times to 28.82 times (Figure 5D). Therefore, the transfer of heterologous or native disulfide bond formation modules by KmYACs increased the overall expressions of disulfide bond formation genes, likely enhancing the process of disulfide bond formation and facilitating secretory expression.

**Figure 5.**
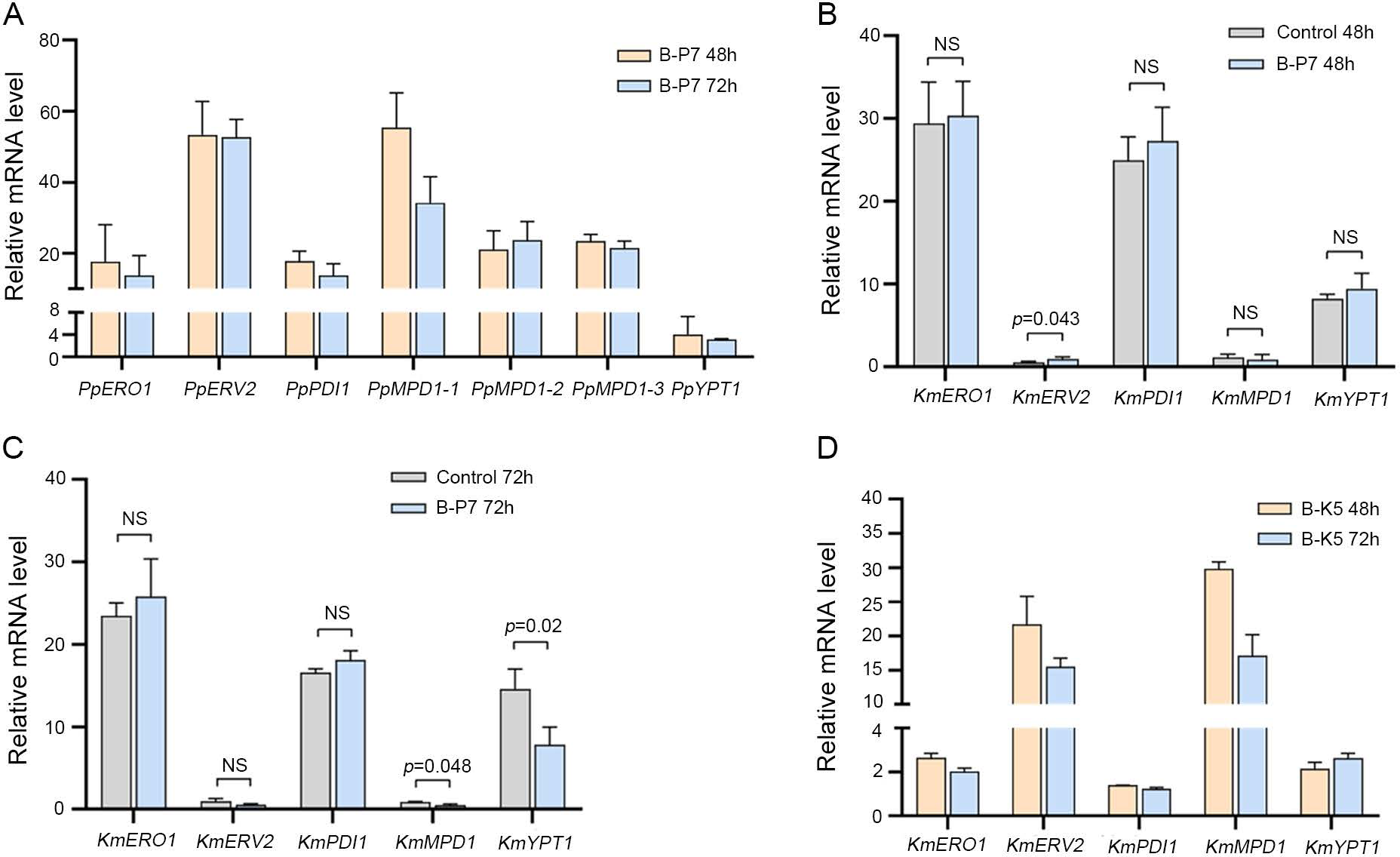
Expressions of the modules genes in B-P7 and B-K5. (A) Relative mRNA levels of P7 module genes in B-P7. B-P7 cells were collected after cultivating 48 h and 72 h. mRNA levels of P7 module genes were normalized to that of *SWC4*. Values represent mean ± S.D. (n=3). (B-C) Relative mRNA levels of *K. marxianus* genes homologuous to P7 module genes. Cells that did not contain KmYACs were used as a control. B-P7 and control cells were collected after cultivating 48 h (B) and 72 h (C). The mRNA levels of homologous genes in B-P7 and control cells were normalized to that of *SWC4*, and the relative levels were compared side by side. NS represents *p*>0.05. (D) Relative mRNA levels of K5 module genes in B-K5. B-K5 cells were collected after cultivating 48 h and 72 h. mRNA levels of K5 module genes in B-K5 were normalized to those of the same genes in the controls cells.

### Transcriptomic and metabolomic analysis of cells containing KmYACs

To get an insight into the mechanism underlying the improved yield of heterologous proteins in cells containing KmYACs, we analyzed the transcriptome and metabolome of B-P7 and B-K5, and compared them with control cells (Figure 6A). We identified a total of 1406 differentially expressed genes (DEGs) between B-P7 and control cells, as well as 2068 DEGs between B-K5 and control cells. A total of 939 DEGs were found to be shared by B-P7 and B-K5 (Table S6). Subsequently, the identified DEGs underwent GO term enrichment analysis. Among the shared upregulated genes, genes implicated in mitochondrial translation, ergosterol biosynthetic process, ubiquinone biosynthetic process, and vesicle-mediated transport were significantly enriched. Genes implicated in DNA integration, RNA-dependent DNA biosynthetic process, and vitamin B6 catabolic process were significantly enriched among the shared downregulation genes (*p*<0.05, Figure 6A). Untargeted metabolomics analysis revealed that 268 and 308 metabolites were altered in B-P7 and B-K5, respectively, compared with control cells (*p*<0.1). Pathways of pyrimidine metabolism, purine metabolism, pentose phosphate pathway, lysine biosynthesis, nicotinate and nicotinamide metabolism, and TCA cycle were both altered in B-P7 and B-K5 (Figure 6B-C).

**Figure 6.**
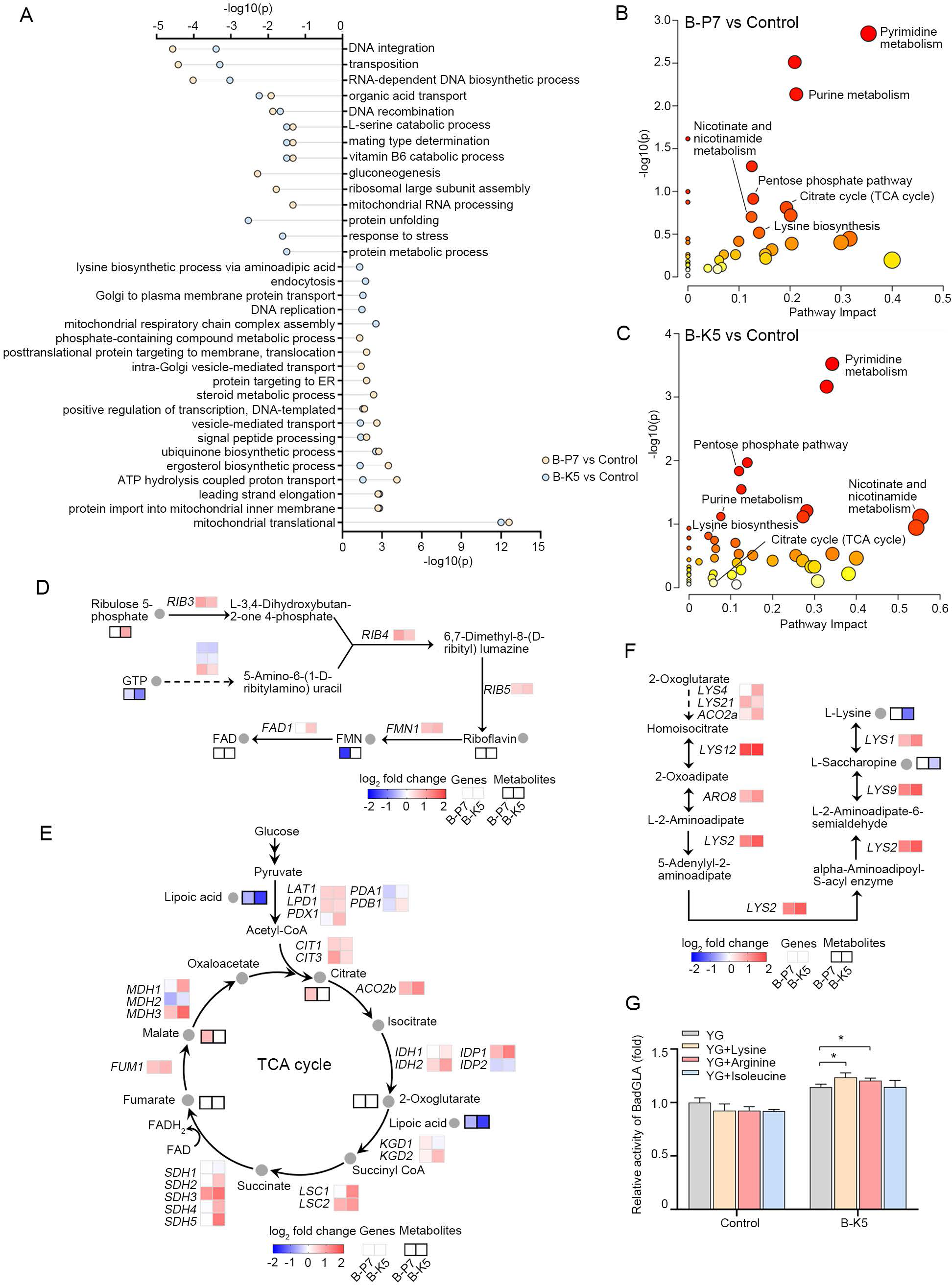
Transcriptomic and metabolomic analysis of B-P7 and B-K5. (A) GO term enrichment analysis of differentially expressed genes (DEGs). Cells that did not contain KmYACs were used as a control. DEGs were identified between B7 cells and control cells, as well as between B-K5 cells and control cells (|Fold Change|≥1.5). DEGs were subjected to GO enrichment analysis. The *p*-values of enrichment are shown on the x-axis (*p*<0.05). (B-C) Metabolomic analysis of B-P7 and B-K5. Changes in metabolites were identified between B-P7 and control cells (B), as well as between B-K5 cells and control cells (C) (*p*<0.1). Metabolic pathway enrichment was performed using MetaboAnalyst, based on the changes in metabolites. The *p*-value of the enrichment and pathway impact are shown in the graph. (D-F) Changes in gene expressions and metabolites in the FAD biosynthesis (D), TCA cycle (E) and lysine biogenesis (F). Levels of gene expressions in B-P7 or B-K5 were compared to those in control cells (*p*<0.05), as well as levels of metabolites were also compared (*p*<0.1). Gray-bordered boxes are used to indicate changes in gene expressions, whereas black-bordered boxes are used to mark changes in metabolites. Metabolites that did not exhibit any significant change are labelled as blank in the boxes. (G) Secretory activity of BadGLA expressed by B-K5 and control cells in YG medium supplemented with 2 mM lysine, arginine or isoleucine. Values represent mean ± S.D. (n=3) (* *p*<0.05).

Consistent changes in gene expression and metabolites in the same pathway suggest that the pathway is significantly altered with a high degree of confidence. For instance, six out of eight genes required for FAD biosynthesis were upregulated in B-P7 and B-K5, while the amount of precursor FMN and GTP was reduced in both strains (Figure 6D). These findings suggest that enhanced disulfide bond formation modules are associated with enhanced FAD biosynthesis, which is expected given that FAD functions as a cofactor of Ero1 and Erv2 in the oxidation of Pdi1^37^.

The expression of pyruvate dehydrogenase complex genes, specifically *LAT1*, *LPD1*, and *PDX1*, was upregulated in both B-P7 and B-K5 strains. Additionally, the amount of lipoic acid, which serves as a substrate of pyruvate dehydrogenase, was consistently reduced in both strains. These results suggest that there is a preferred metabolic flux from pyruvate into acetyl-CoA. Following this, the expressions of citrate synthase (Cit1 and Cit3), which catalyze the production of citrate from acetyl-CoA, were upregulated, indicating an enhanced flux entering the TCA cycle. Most of the enzymes in the TCA cycle were upregulated in both strains, with the upregulation being more significant in B-K5. Therefore, it seems that increased flux into the TCA cycle is a feature of cells overexpressing heterologous proteins (Figure 6E).

Amino acids serve as building blocks for protein biosynthesis. In B-K5, all eight genes required for lysine biogenesis displayed unanimous upregulation, while the amounts of lysine and its precursor, saccharopine, were reduced (Figure 6F). Notably, this unanimous upregulation of synthetic genes and reduction of the end product was not observed in biogenesis pathways of other amino acids. Therefore, these results strongly indicate a high demand for lysine in B-K5. To investigate this further, we supplemented the medium of B-K5 with 2 mM lysine and examined its effect on the yield of heterologous proteins. Compared to the medium without supplementation, the addition of lysine resulted in an 8% increase in secretory activities of BadGLA (Figure 6G). Arginine has similar characteristics to lysine, as both are basic amino acids. Six out of seven genes required for arginine biogenesis were upregulated in B-K5, including the rate-limiting gene *ARG1* (Figure S4). Same to the effects of adding lysine, adding arginine increased secretory activities by 5.3% (Figure 6G). As a control, adding isoleucine did not improve the yield. This was consistent with the transcriptomic results, as *CHA1*, the rate-limiting gene of isoleucine biogenesis, was significantly downregulated in B-K5 (Figure S4). Therefore, cells overexpressing heterologous proteins might have a preferred demand for lysine and arginine.

## Discussion

In this study, YAC was constructed for *K. marxianus* for the first time. The effects of telomeres (*Tetrahymena* or *K. marxianus*) and configuration (linear or circular) on the stability of KmYAC were evaluated. Telomeres from *Tetrahymena* supported the stable propagation of KmYACs as well as *K. marxianus* telomeres, indicating functional telomeres were formed at the end of *Tetrahymena* telomeres. It was observed that yeast can recognize and use telomeres from distantly related organisms. During the replication of linear plasmids in *S. cerevisiae*, telomere repeat units of *S. cerevisiae* C_1-3_A were added to the ends of *Tetrahymena* telomeres C_4_A_2_^41^. Our results suggest that the exceptionally long telomeric repeat unit of *K. marxianus* (25 bp) can also be added to the ends of *Tetrahymena* telomeres containing short repeat units (6 bp). Therefore, it may be possible to construct YACs compatible with yeast species having long telomeric repeat units, such as *Candida maltosa*, *Candida pseudotropicalis*, *Candida tropicalis*, *K. lactis*, and *S. kluyveri* by using heterologous telomeres with short repeat units (6-8 bp)^42–43^. Regarding the effect of configuration, our results showed that there was no difference in stability between circular and linear KmYACs. Moreover, both types of KmYACs did not affect the growth of the host. These results are consistent with a previous report on *S. cerevisiae*, where both linear and circular neochromosomes were present in one copy per cell, stable, and innocuous to their host^44^. Circular YACs up to 600 kb can be purified from natural chromosomes for further manipulation, exhibiting better operability compared to linear YACs^45^. Therefore, circular YACs can serve as an alternative to linear YACs for transferring large functional modules in both *S. cerevisiae* and unconventional yeasts.

The largest module loaded into KmYACs was 15 kb, well below the maximum capacity of YAC to load exogenous DNA (∼3 Mb). This was primarily because the disulfide formation modules contained rationally designed promoters, ORFs, and terminators. The rational design approach limited the size of a single fragment obtained either by chemical synthesis or PCR amplification. To address this issue, we used Gibson assembly to ligate two smaller cassettes, instead of a single long cassette, with the KmYAC shuttle vector in vitro (Figure 1). To fully utilize the loading capacity of KmYAC in the future, more cassettes might be assembled with the KmYAC vector in vivo, using transformation-associated recombination (TAR). A similar strategy was successfully applied in *K. marxianus*, where five fragments were ligated together^13^. Meanwhile, random recombination between cassettes could be performed to construct diverse modules in KmYACs, resulting in a library for screening purposes. For example, seven enzymes from the flavonoid pathway were individually cloned into expression cassettes and then randomly combined on YACs to screen for specific flavonoid compounds in *S. cerevisiae*^27^. Therefore, by introducing novel strategies to improve the loading capacity and diversity of assemblies, KmYACs are expected to find more applications in metabolic engineering and synthetic biology.

Although *K. marxianus* is not as widely used as *P. pastoris* in the production of heterologous proteins, it is regarded as a promising cell factory due to its various advantageous features. For example, *K. marxianus* has been reported as the fastest-growing eukaryote, with a specific growth rate of up to 0.80 h^−1^ ^46^. The fermentation of *K. marxianus* usually takes 2-3 days^5^. In contrast, *P. pastoris* exhibits a specific growth rate of only 0.18 h^−1^, and it usually takes 7-8 days to complete fermentation^47^. *K. marxianus* is well-known for its thermotolerance and its production performance remains stable between 30-37°C^2, 40^. On the other hand, *P. pastoris* requires a growth temperature of 28∼30 °C, and its production performance rapidly deteriorates at 32°C^48^. Furthermore, the expression of heterologous proteins in *P. pastoris* relies on methanol induction, which may pose risks to both the production process and the safety of the products^48^. In contrast, *K. marxianus* uses constitutive promoters to express proteins and its culture medium contains inorganic salts and glucose, ensuring the safety of both production and products^5^. However, the maximum yield of heterologous proteins in *K. marxianus* remains lower than that in *P. pastoris*. Therefore, transferring essential genes from *P. pastoris* that control protein folding and maturation into *K. marxianus* is a feasible strategy to enhance protein yield. Given this consideration, five or seven genes from the disulfide bond formation modules of *P. pastoris* were introduced into *K. marxianus* using KmYAC. These genes effectively functioned in vivo, and their expressions were compatible with the genes involved in the native disulfide bond isomerization pathway in *K. marxianus* (Figure 5).

Transcriptomic and metabolomic analyses showed that the introduction of disulfide bond formation modules via KmYAC caused significant changes in various cellular processes, which may have contributed to the improved yield of heterologous protein in B-P7 and B-K5 (Figure S5). Among the enhanced processes, the TCA cycle, mitochondrial respiratory chain, ergosterol synthesis, and DNA synthesis were also upregulated in *S. cerevisiae* cells exhibiting a high yield of heterologous proteins^49–51^, suggesting that upregulation of these processes might be a conserved response to the stress caused by overexpression of heterologous proteins (Figure S5). While the upregulation of FAD synthesis suggests active oxidation of disulfide bonds inside ER, upregulation of the thioredoxin and glutathione antioxidant systems might be required to counteract the aberrant oxidation of free sulfhydryl groups in the intracellular environment (Figure S5)^37, 52^.

Adjusting the amount of a single amino acid in the medium has different effects on the yield of heterologous proteins. For instance, when the amount of methionine in the medium was reduced, the expression of amyloid-β peptides in *S. cerevisiae* increased by 2 folds. On the other hand, increasing the concentration of cystine in the medium did not affect the yield^51^. In this study, transcriptomic and metabolomic analysis of B-K5 suggests an urgent demand for the supply of arginine and lysine, but not for isoleucine. Consistently with these findings, adding lysine or arginine, but not isoleucine, further improved the yield of a heterologous protein in the B-K5. Lysine and arginine are basic amino acids commonly found in the target sequence of mitochondrial proteins^53–54^. The import of mitochondrial proteins was significantly upregulated in B-K5 (Figure 6A), and supplementation with arginine and lysine might improve this process, which could be coupled with the enhanced mitochondrial respiratory chain in B-K5. Furthermore, the integration of vital cofactors into proteins, such as the covalent attachment of lipoic acid in the E2-subunit of the pyruvate dehydrogenase complex, is made possible by the ε-amino group of lysine^54^. Therefore, supplementation with lysine might promote the production of acetyl-CoA, which was consistent with a preferred metabolic flux into the TCA cycle in B-K5 (Figure 6E). In further studies, it would be intriguing to investigate the effect of supplementing other amino acids on the yield.

## Materials and methods

### Strains and medium

A *K. marxianus* FIM1 strain in its wild type was deposited at China General Microbiological Culture Collection Center (CGMCC, No 10621)^5^. *URA3*, *HIS3*, and *TRP1* were deleted in FIM1 and the resultant strain was named FIM-1ΔUΔHΔT, which was used as a host strain in this study^55^. Cells were cultivated at 30 ℃. FIM1 and FIM-1ΔUΔHΔT were grown in YPD medium (2% peptone, 1% yeast extract, 2% agar for plates). Synthetic dropout medium without histidine and tryptophan (SC-His-Trp) and that without uracil (SC-Ura) were prepared as described before^56^. For expressing heterologous proteins, cells were grown in a YG medium (2% yeast extract, 4% glucose).

### Plasmids

Plasmids and primers used in the study are listed in Supplementary Table S1. KmYAC shuttle vectors were synthesized by Genewiz (Nanjing, China). KmYAC was composed of *HIS3*, *Escherichia coli* origin (OriC), ampicillin-resistant gene (*Amp^R^*), *ARS1*/*CEN5*, *HphMX4*, *TRP1* and a TEL-filler-TEL cassette. *ARS1/CEN5*, *HIS3* and *TRP1* originate from the genome of FIM1. *HphMX4* originates from pCloneHyg1^57^. A *Not* I site was inserted into the open reading frames (ORF) of *HphMX4* without changing the amino acid sequence. OriC and *Amp^R^* originate from pUC57 vector (Genewiz). The TEL-filler-TEL cassette contained two inverted telomeres (*Tetrahymena* or *K. marxianus*) separated by a filler sequence and two flanking *BamH* I sites. Telomere contains 44 tandem repeats of *Tetrahymena* telomeric sequence (5’-GGGGTT-3’) or 25 tandem repeats of *K. marxianus* telomeric sequence (5’-GGTGTACGGA TTTGATTAGT TATGT-3’). KmYAC shuttle vector containing *Tetrahymena* and *K. marxianus* telomeres was named LHZ1014 and LHZ1015, respectively. Sequences of LHZ1014 and LHZ1015 are listed in Supplementary Table S2-S3. ORFs of *PpERO1* (XM_002489600), *PpERV2* (XM_002492510), *PpPDI1* (XM_002494247), *PpMPD1-1* (XM_002489421), *PpMPD1-2* (XM_002489761), *PpMPD1-3* (XM_002494173) and *PpYPT1* (XM_002492399) were amplified from the genome of *P. pastoris* GS115, as well as those of *KmERO1* (XM_022818302), *KmERV2* (XM_022821364), *KmPDI1* (XM_022822028), *KmMPD1* (XM_022822290), *KmYPT1* (XM_022821297) were amplified from the genome of FIM1-ΔU. Strong promoters, including P*_ENO2_*, P*_PDC1_*, P*_INU1_*, P*_HXT4_*, P*_OM45_*of *K. marxianus*, P*_AgTEF_* of *Ashbya gossypii*, P*_ScADH1_* of *S. cerevsiae*, and synthetic terminators, including T*_synth3_*, T*_synth8_*, T*_synth19_*, T*_synth27_*, T*_synth28_*, T*_synth30_* were described previously^58–59^. The *PpPDI1* cassette (P*_INU1_*-*PpPDI1*-T*_synth28_*), the *PpMPD1-2* cassette (P*_AgTEF_*-*PpMPD1-2*-T*_synth30_*), the *PpYPT1* cassette (P*_OM45_*-*PpYPT1*-T*_synth8_*), and the *PpERV2* cassette (P*_PDC1_*-*PpERV2*-T*_synth19_*) were ligated in tandem with pMD18-T (Takara, Beijing, China) to construct LHZ1016. The *PpMPD1-1* cassette (P*_HXT4_*-*PpMPD1-1*-T*_synth19_*), the *PpMPD1-3* cassette (P*_ScADH1_*-*PpMPD1-3*-T*_synth3_*), and the *PpERO1* cassette (P*_ENO2_*-*PpERO1*-T*_synth27_*) were ligated with pMD18-T to construct LHZ1017. The *KmPDI1* cassette (P*_INU1_*-*KmPDI1*-T*_synth28_*), the *KmMPD1* cassette (P*_HXT4_*-*KmMPD1*-T*_synth19_*), and the *KmYPT1* cassette (P*_OM45_*-*KmYPT1*-T*_synth8_*) were ligated with pMD18-T to construct LHZ1018. The *KmERV2* cassette (P*_PDC1_*-*KmERV2*-T*_synth19_*) and the *KmERO1* cassette (P*_ENO2_*-*KmERO1*-T*_synth27_*) were ligated with pMD18-T to construct LHZ1019. *BadGLA* from *Blastobotrys adeninivorans* was codon-optimized for *K. marxianus* and inserted between *Spe* Ⅰ and *Not* Ⅰ sites of pUKDN132 to obtain LHZ1020^5,60^. *TeGlaA* from *Talaromyces emersonii* was codon-optimized for *K. marxianus* and inserted between *Spe* Ⅰ and *Not* Ⅰ sites of pUKDN132 to obtain LHZ1021. LHZ745 expressing *Xyn-CDBFV* was described previously^59^.

### Construction of the yeast artificial chromosome

The tandem cassette containing *PpPDI1*, *PpMPD1-2*, *and PpYPT1* was amplified from LHZ1016 and named P5 cassette Ⅰ, and the cassette containing *PpMPD1-1*, *PpMPD1-3* was amplified from LHZ1017 and named P5 cassette Ⅱ. The tandem cassette containing *PpPDI1*, *PpMPD1-2*, *PpYPT1*, and *PpERV2* was amplified from LHZ1016 and named P7 cassette Ⅰ, and the cassette containing *PpMPD1-1*, *PpMPD1-3*, *PpERO1* was amplified from LHZ1017 and named P7 cassette Ⅱ. The tandem cassette containing *KmPDI1*, *KmMPD1*, and *KmYPT1* was amplified from LHZ1018 and named K5 cassette Ⅰ, and the cassette containing *KmERV2*, *KmERO1* was amplified from LHZ1019 and named K5 cassette Ⅱ. To construct the linear KmYAC, LHZ1014 or LHZ1015 was digested with *Not* Ⅰ and *BamH* Ⅰ to release two arms and the filler sequence. After dephosphorylation, the digested product of LHZ1014 was ligated with P5 cassette I and II by Gibson assembly to form Lin Te_TEL-P5, ligated with P7 cassette I and II to form Lin Te_TEL-P7, and ligated with K5 cassette I and II to form Lin Te_TEL-K7. Similarly, the digested product of LHZ1015 was ligated with P5 cassettes to form Lin Km_TEL-P5, ligated with P7 cassettes to form Lin Km_TEL-P7, and ligated with K5 cassettes to form Lin Km_TEL-K5. To construct the circular KmYAC, LHZ1015 was digested with *Not* Ⅰ and dephosphorylated. The digested product was ligated with P5 cassettes to form Cir Km_TEL-P5, ligated with P7 cassettes to form Cir Km_TEL-P7, and ligated with K5 cassettes to form Cir Km_TEL-K5. The ligation product was transformed into FIM-1ΔUΔHΔT by lithium acetate method^61^, and selected on SC-His-Trp plates. Transformants were identified by colony PCR. Primers used in the construction are listed in Supplementary Table S1.

### Enzymatic assays and SDS-PAGE

LHZ1020, LHZ1021 or LHZ745 was transformed into cells containing KmYACs. Transformants were selected on SC-Ura plates and cultivated in 50 mL YG medium at 30 ℃ for 72 h. The supernatant was collected for enzymatic assays and SDS-PAGE analysis. The activities of glucoamylases (BadGLA and TeGlaA) were determined by using 1% (w/v) soluble starch as a substrate. The activity of Xyn-CDBFV was determined by using 1% wheat arabinoxylan as a substrate. One unit of activity of these three enzymes was defined as the amount of enzyme required to release 1 μmol glucose per minute^59^. Fed-batch fermentations of strains expressing BadGLA were conducted in a 5-L bioreactor (BXBIO, Shanghai, China) as described previously^5^. The supernatant was collected after 24, 48, 60 and 72 h. Samples were subjected to the SDS-PAGE analysis and enzymatic assay.

### Southern blot

Southern blot was performed as described before^62^. In brief, cells containing KmYACs were embedded in agarose plugs. Spheroplasts were created and lysed in the plugs. Subsequently, the plugs were subjected to the pulsed field electrophoresis (Bio-Rad, California, USA). Linear KmYACs were resolved directly. Circular KmYACs were entrapped in the agarose plugs and separated from the natural chromosomes during electrophoresis. Plugs containing circular KmYACs were digested with *Not* I and subjected to the pulsed-field electrophoresis again to release the linearized KmYACs. KmYACs were transferred from gel to a nylon membrane (Beyotime, Shanghai, China) for about 16-20 h. The membrane was hybridized with DNA probes prepared using a Biotin Random Primer DNA Labeling Kit (Beyotime, Shanghai, China) and then detected using Chemiluminescent Biotin-labeled Nucleic Acid Detection Kit (Beyotime, Shanghai, China). The probe targets *HphMX4* and its sequence was listed in Supplementary Table S1.

### Determination of copy numbers of KmYACs

Transformants containing KmYACs were grown in a 3 mL SC-His-Trp medium overnight. Genomic DNA was extracted from the cells and subjected to quantitative PCR, as described previously^59^. The copy number of KmYACs was determined by comparing the level of *HphMX4* to that of endogenous *LEU2*^59^.

### Stabilities of KmYACs

Transformants containing KmYACs were cultivated in the 3 mL SC-His-Trp medium overnight. The resulting start culture was diluted into 50 mL YPD medium at an OD_600_ of 0.2 and grown for 24 h (about 7 generations). This process was repeated for the next 4 days, resulting in a total of 35 generations. Cells from both the initial culture and final culture were collected and spread onto YPD or SC-His-Trp plate at an appropriate dilution. The stability of the plasmid was determined as the ratio of colonies formed on the SC-His-Trp plate to that on the YPD plate. The loss percentage of the KmYAC per generation was calculated as described previously^15^. The experiment was performed with three parrallel culture.

### Growth curves of cells containing KmYACs

Cells containing KmYACs were grown in 3 mL SC-His-Trp medium. The culture was diluted into 50 mL YPD medium at an OD_600_ of 0.2 and grown for 72 h at 30℃. OD_600_ of the culture was measured after 6, 9, 12, 24, 30, 36, 48, 54, 60 and 72 h. The experiment was performed with three parrallel culture.

### qPCR and RNA-seq

Transformants containing LHZ906 and KmYAC (Lin Km_TEL-P7 or Lin Km_TEL-K5) were cultivated in the 3 mL SC-His-Trp medium at 30℃ overnight. The overnight cultures were diluted into 50 mL YG medium at an OD_600_ of 0.2 and grown at 30℃ for 72 h. Cells were collected at 48 h and 72 h after cultivation. For qPCR, RNA was extracted from frozen cells using Quick-RNA Fungal/Bacterial Miniprep kit (Zymo Research, California, USA) and was reverse transcribed using a PrimeScript RT Reagent Kit (Takara, Beijing, China). qPCR was performed using TB Green Premix Ex Taq (Takara, Beijing, China). Primers used in qPCR are listed in Supplementary Table S1. For RNA-seq, RNA was extracted from cells collected at 48 h by using TRIzol Reagent (Invitrogen, California, USA), reversed transcribed using TruSeqTM RNA sample preparation Kit (Illumina, California, USA) and sequenced by Illumina NovaSeq 6000 system (BIOZERON, Shanghai, China). After the quality control, the raw paired-end reads (150 bp*2) were separately aligned to the *K. marxianus* FIM1 reference genome and preprocessed by the RNA-seq pipeline from Shanghai BIOZERON Co., Ltd. Differential expression genes (DEGs, |Fold Change|≥1.5) were significantly enriched in GO terms, in which the *p*-value of GO terms was less than 0.05. RNA-seq data is listed in Supplementary Table S4.

### Metabolomics analysis

Cells were collected at 48 h after cultivation as described in RNA-seq. Metabolites were extracted from cells for untargeted metabolomics analysis (BIOZERON, Shanghai, China). UHPLC-MS/MS analyses were conducted utilizing a Vanquish UHPLC system (Thermo Fisher, MA, Germany) connected to an Orbitrap Q Exactive™ HF mass spectrometer (Thermo Fisher, MA, Germany). Metabolic pathway analysis was carried out using MetaboAnalyst (http://www.metaboanalyst.ca). A *p*-value of <0.1 was considered statistically significant for differential metabolites, while a *p*-value of <0.05 was considered statistically significant for metabolic pathway enrichment. Metabolomic data is listed in Supplementary Table S5.

### Statistical analysis

Each assay in the study contained three biological replaicates. Unpaired two tailed t-test was used to calculate the statistical significance. *p*-value< 0.05 was considered as statistically significant.

## Acknowledgements

This study was supported by the National Key Research and Development Program of China 2021YFA0910601 and 2021YFC2100203, Shanghai Municipal Education Commission (2021-03-52) and Science and Technology Research Program of Shanghai 19DZ2282100.

## Author Contributions

**Pingping Wu**: Conceptualization, Data curation, Formal analysis, Investigation, Methodology, Roles/Writing - original draft. **Wenjuan Mo**: Formal analysis, Software. **Tian Tian**: Data curation, Validation. **Kunfeng Song**: Visualization. **Yilin Lyu**: Methodology**. Haiyan Ren**: Methodology. **Jungang Zhou**: Investigation. **Yao Yu**: Conceptualization, Formal analysis, Methodology, Roles/Writing - original draft. **Hong Lu**: Funding acquisition, Project administration, Supervision, Writing - review & editing.

## Ethics statement

No animal or human research was involved in this study.

## Data availability

Data will be made available on request.

## Supplementary Information

**Supplementary Figure S1.**
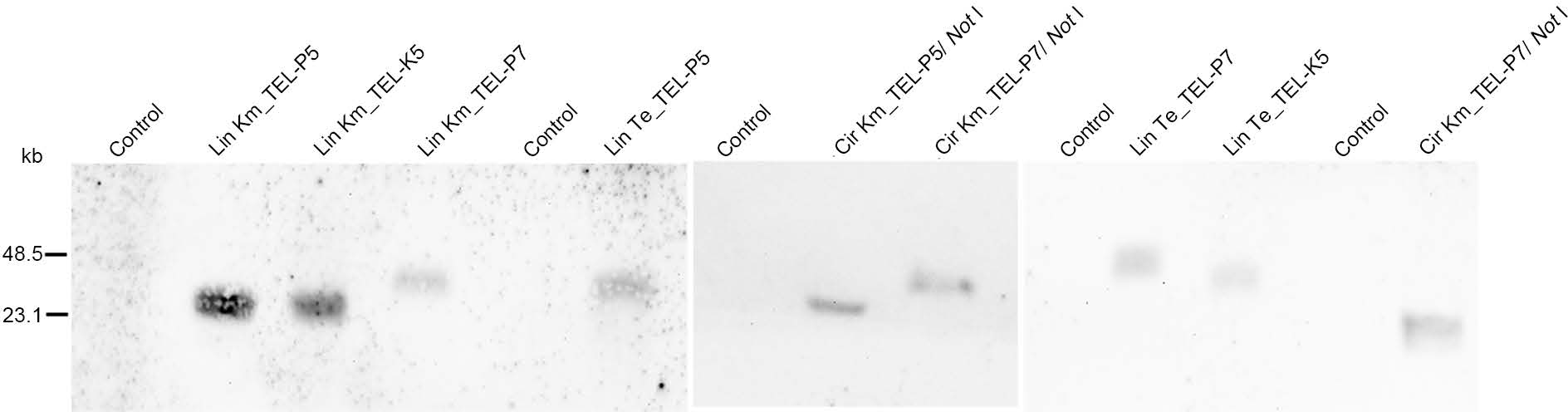
Southern blot of nine KmYACs in the cells. Circular KmYACs were digested with *Not* before blotting.

**Supplementary Figure S2.**
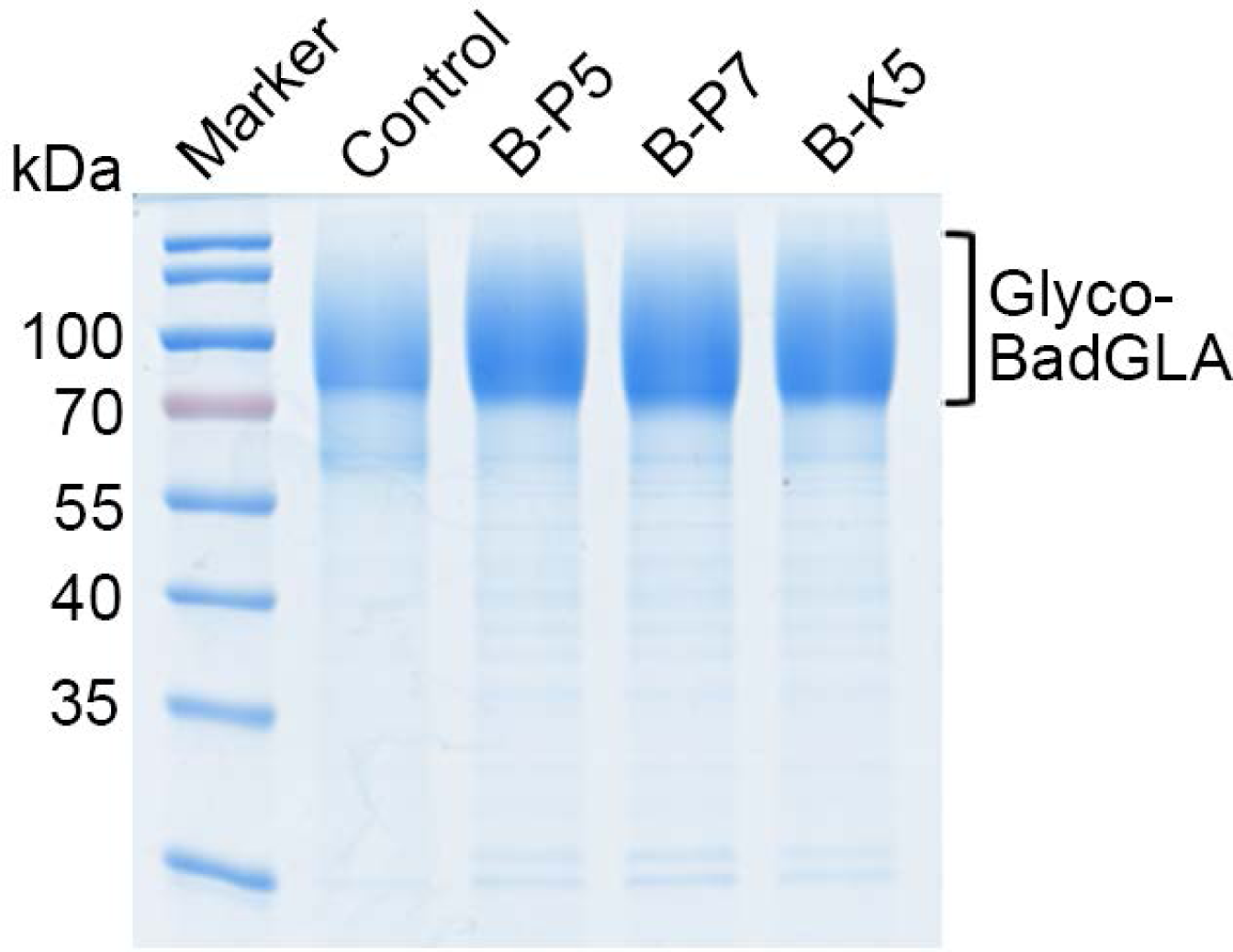
SDS-PAGE of the samples in a 5 L fermenters diluted for 35 times.

**Supplementary Figure S3.**
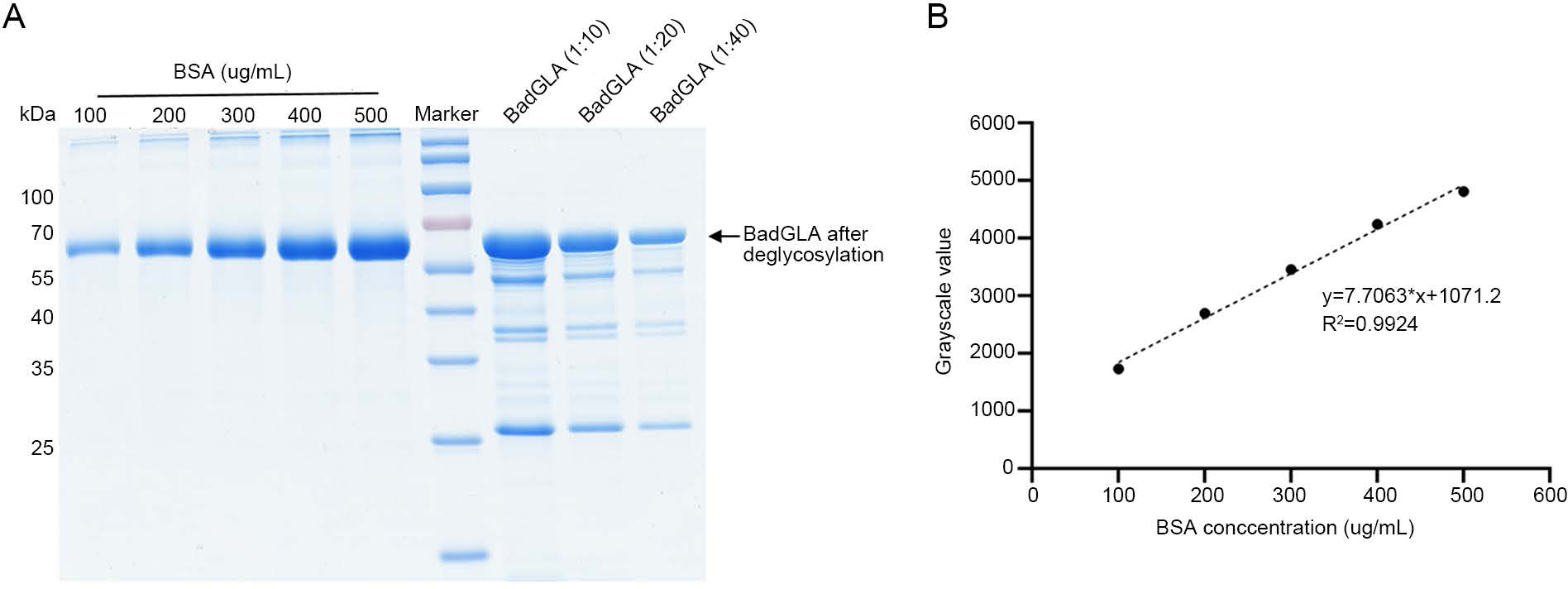
Determination of the specific acitivity of BadGLA. (A) SDS-PAGE of deglycosylated BadGLA. FIM-1ΔU was transformed with LHZ906 and transformatns were grown in YG medium for 72 h. The BadGLA in the supernatant was deglycosylated using Endo H (NEB, MA, USA). Different dilutions of the deglycosylated BadGLA were separated using SDS-PAGE, alongside dilutions of bovine serum albumin (BSA). (B) Linear fitting analysis of BSA concentrations and grayscale values. The SDS-PAGE gel in (A) was scanned, and the concentrations of BSA were fitted with the grayscale values of the bands using a linear equation. Using this equation, the concentration of BadGLA in the sample was determined to be 256.386 ug/mL. Since the activity of the BadGLA sample was 127.685 U/mL, the specific activity of BadGLA was determined to be 498.02 U/mg.

**Supplementary Figure S4.**
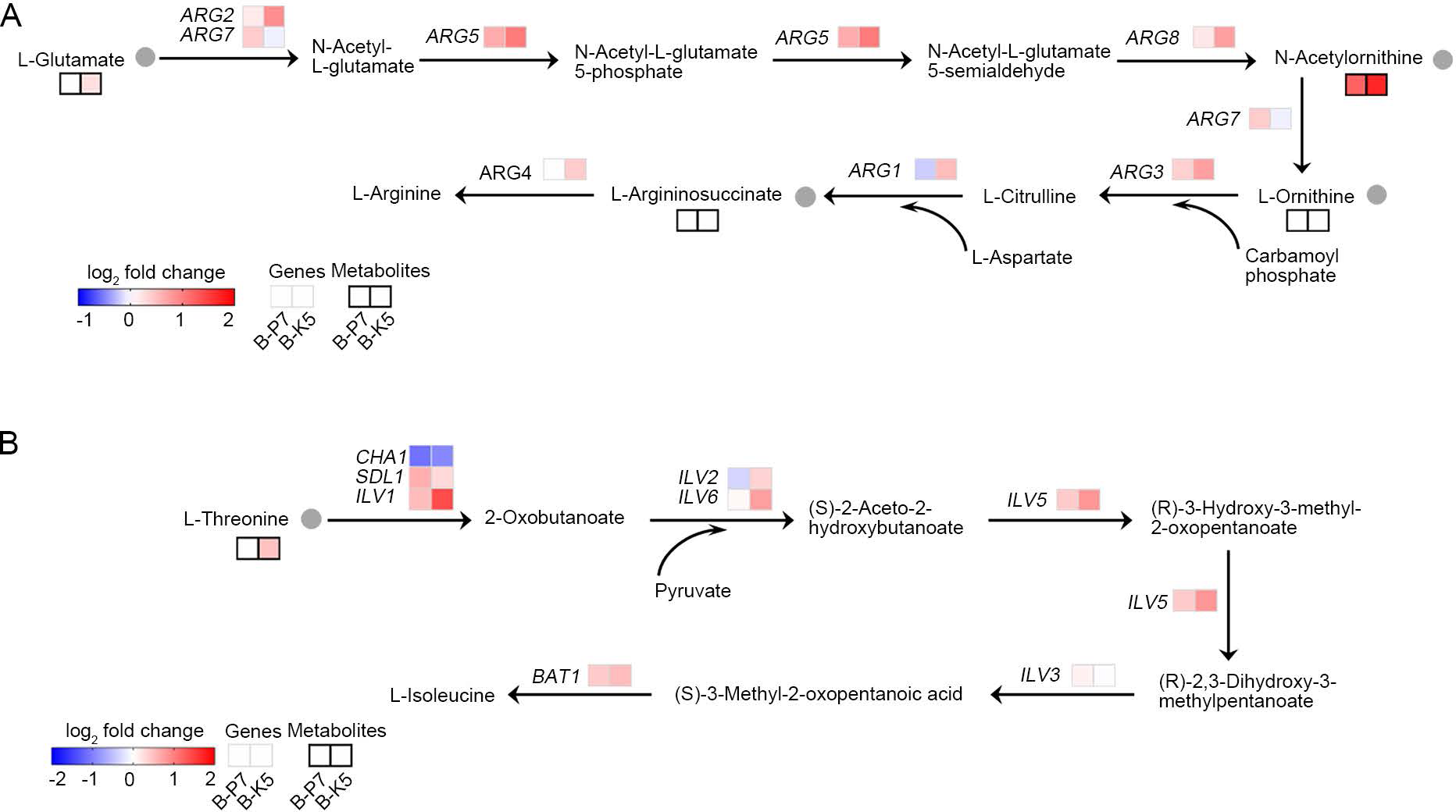
Changes in gene expressions and metabolites in the arginine and isoleucine biosynthesis. (A-B) Expression levels of genes in the arginine (A) and isoleucine biosynthesis (B) in B-P7 or B-K5 were compared to those in control cells (*p*<0.05), as well as levels of metabolites were also compared (*p*<0.1). Gray-bordered boxes are used to indicate changes in gene expressions, whereas black-bordered boxes are used to mark changes in metabolites. Metabolites that did not exhibit any significant change are labelled as blank in the boxes.

**Supplementary Figure S5.**
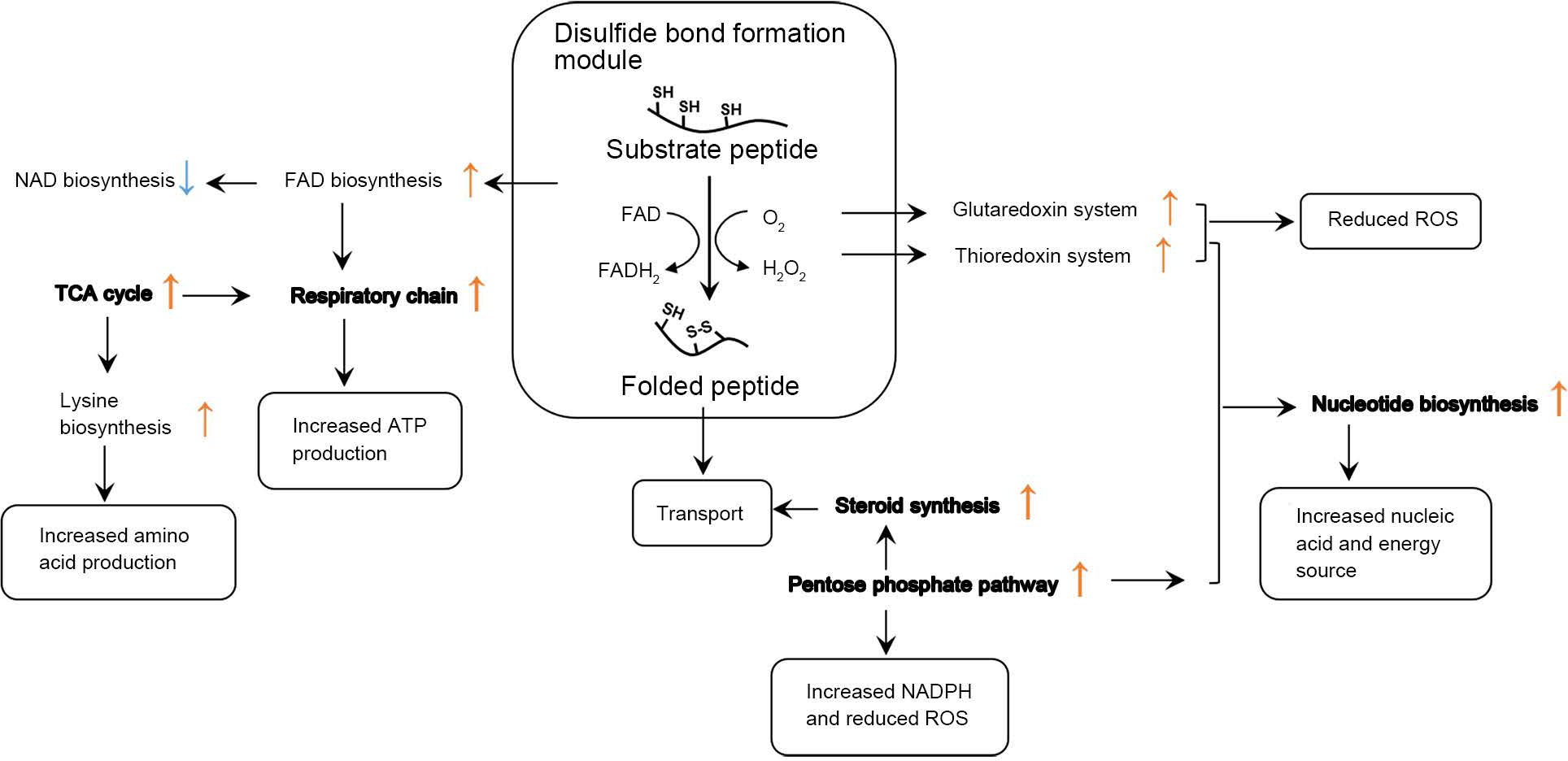
A model describing global changes in diverse processes in B-P7 and B-K5. The figure shows the relationships between these processes and the improved yield of BadGLA. Upregulation of FAD biosynthesis promotes the oxidation of disulfide bonds, while enhanced thioredoxin and glutathione systems, as well as activation of the pentose phosphate pathway, might counteract the aberrant oxidation of free sulfhydryl groups in the intracellular environment. Enhanced lysine biosynthesis provides more building blocks for protein production, and upregulation of the TCA cycle and respiratory chain provides more energy support. Additionally, enhanced nucleotide biosynthesis provides a greater source of nucleic acid and energy, and enhanced ergosterol biosynthesis facilitates vesicle transport. Processes observed in *S. cerevisiae* cells exhibiting a high yield of heterologous proteins are labeled in bold.

**Supplementary Table S1.** Plasmids and primers used in this study.

**Supplementary Table S2.** Sequences of LHZ1014.

**Supplementary Table S3.** Sequences of LHZ1015.

**Supplementary Table S4.** RNA-seq data of B-P7 and B-K5.

**Supplementary Table S5.** Metabolomic data of B-P7 and B-K5.

**Supplementary Table S6.** Differentially expressed genes shared by B-P7 and B-K5.

