## Supplementary materials for "Transfer of disulfide bond formation modules via yeast artificial chromosomes promotes the expression of heterologous proteins in *Kluyveromyces marxianus*"

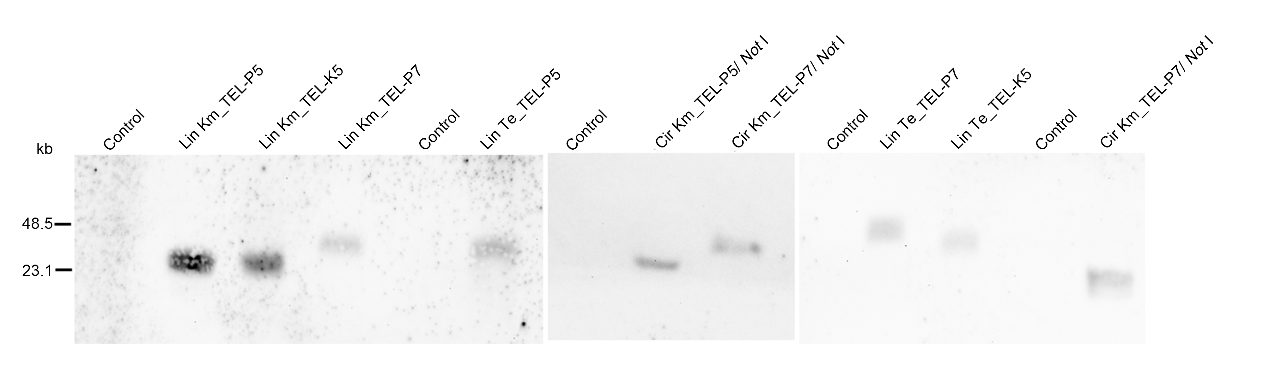

**Supplementary Figure S1**. Southern blot of nine KmYACs in the cells. Circular KmYACs were digested with *Not* Ⅰ before blotting.

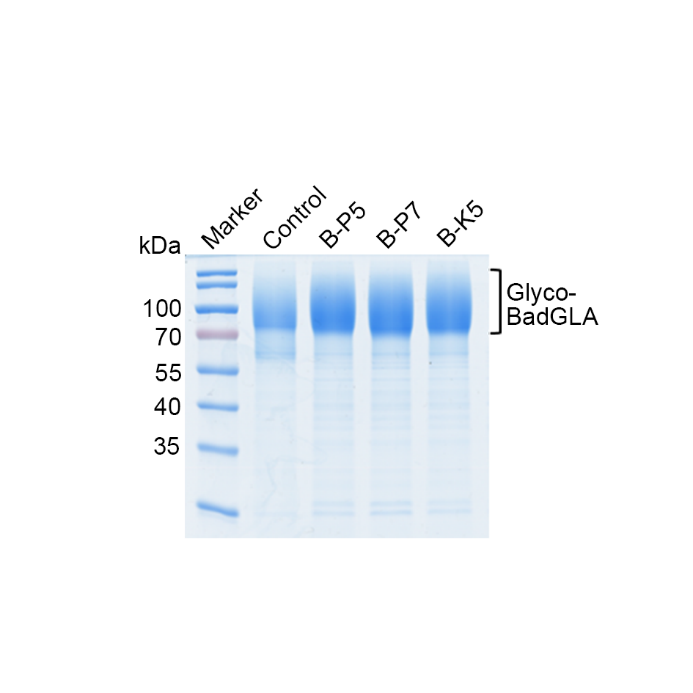

**Supplementary Figure S2**. SDS-PAGE of the samples in a 5 L fermenters diluted for 35 times.

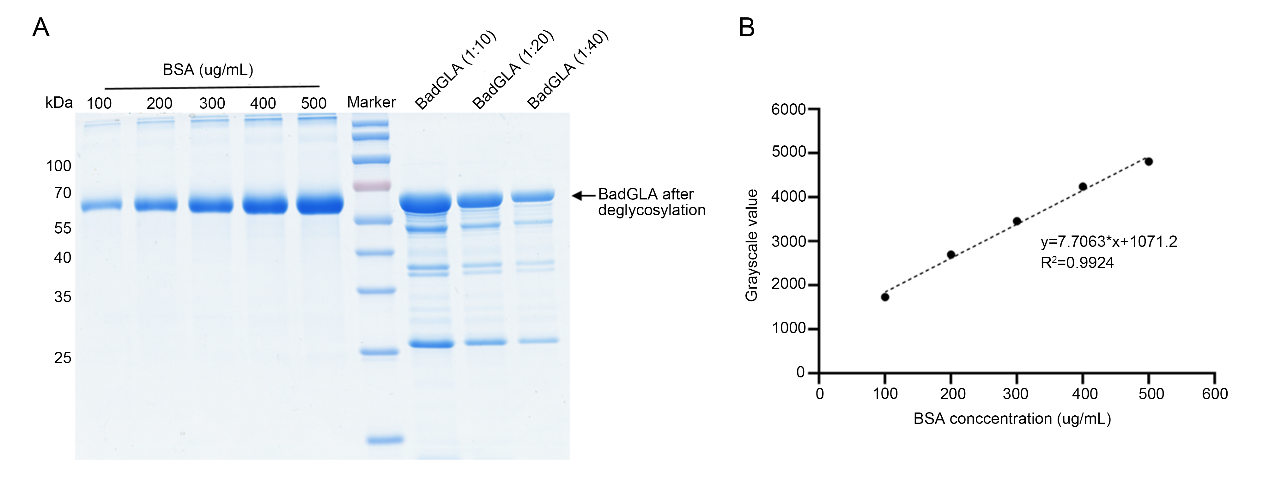

**Supplementary Figure S3**. Determination of the specific acitivity of BadGLA. (A) SDS-PAGE of deglycosylated BadGLA. FIM-1ΔU was transformed with LHZ906 and transformatns were grown in YG medium for 72 h. The BadGLA in the supernatant was deglycosylated using Endo H (NEB, MA, USA). Different dilutions of the deglycosylated BadGLA were separated using SDS-PAGE, alongside dilutions of bovine serum albumin (BSA). (B) Linear fitting analysis of BSA concentrations and grayscale values. The SDS-PAGE gel in (A) was scanned, and the concentrations of BSA were fitted with the grayscale values of the bands using a linear equation. Using this equation, the concentration of BadGLA in the sample was determined to be 256.386 ug/mL. Since the activity of the BadGLA sample was 127.685 U/mL, the specific activity of BadGLA was determined to be 498.02 U/mg.

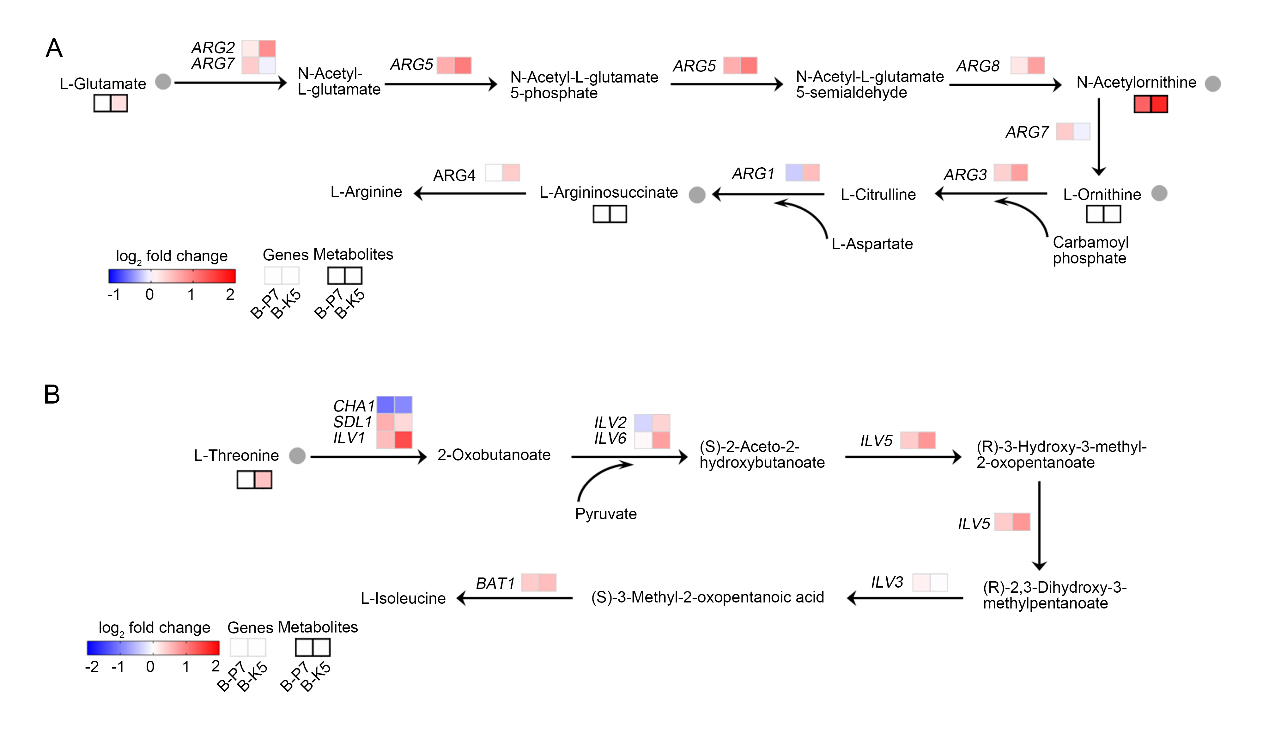

**Supplementary Figure S4.** Changes in gene expressions and metabolites in the arginine and isoleucine biosynthesis. (A-B) Expression levels of genes in the arginine (A) and isoleucine biosynthesis (B) in B-P7 or B-K5 were compared to those in control cells (*p*<0.05), as well as levels of metabolites were also compared (*p*<0.1). Gray-bordered boxes are used to indicate changes in gene expressions, whereas black-bordered boxes are used to mark changes in metabolites. Metabolites that did not exhibit any significant change are labelled as blank in the boxes.

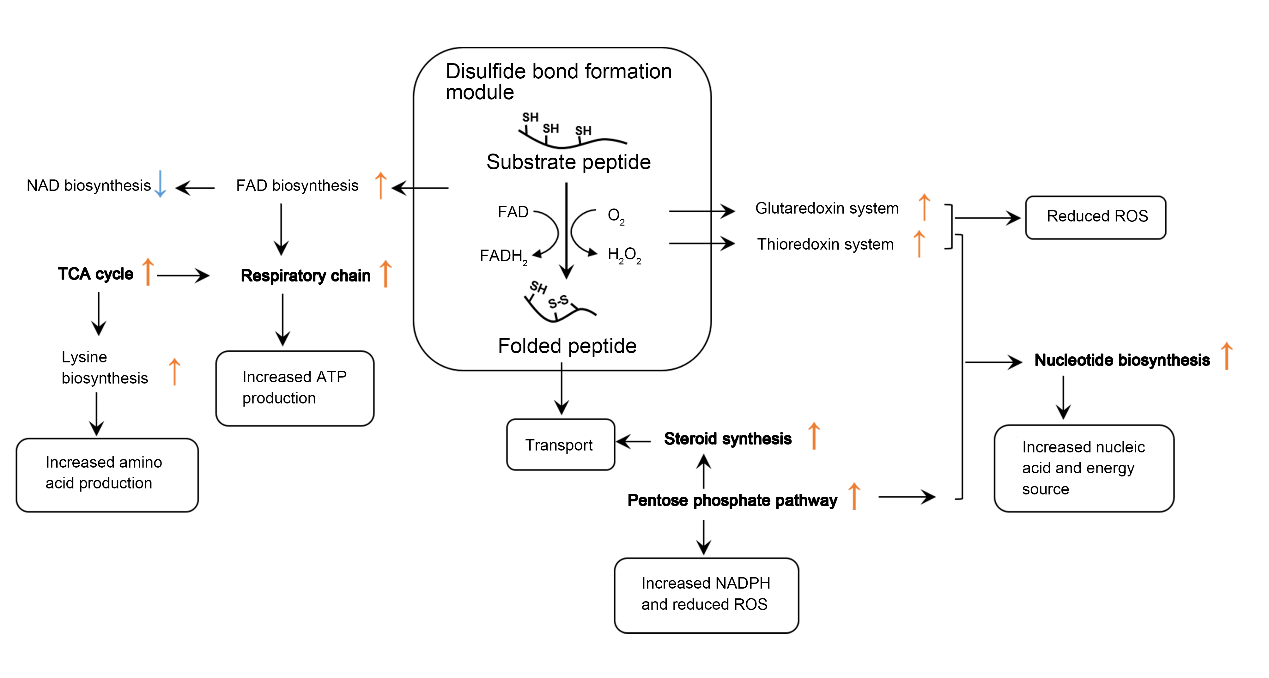

**Supplementary Figure S5.** A model describing global changes in diverse processes in B-P7 and B-K5. The figure shows the relationships between these processes and the improved yield of BadGLA. Upregulation of FAD biosynthesis promotes the oxidation of disulfide bonds, while enhanced thioredoxin and glutathione systems, as well as activation of the pentose phosphate pathway, might counteract the aberrant oxidation of free sulfhydryl groups in the intracellular environment. Enhanced lysine biosynthesis provides more building blocks for protein production, and upregulation of the TCA cycle and respiratory chain provides more energy support. Additionally, enhanced nucleotide biosynthesis provides a greater source of nucleic acid and energy, and enhanced ergosterol biosynthesis facilitates vesicle transport. Processes observed in *S. cerevisiae* cells exhibiting a high yield of heterologous proteins are labeled in bold.
