## Supplementary Table S2 for "Transfer of disulfide bond formation modules via yeast artificial chromosomes promotes the expression of heterologous proteins in *Kluyveromyces marxianus*"

LOCUS	LHZ1014	11687 bp	DNA	circular	10-SEP-2020

SOURCE

ORGANISM

COMMENT	This file is created by Vector NTI

http://www.invitrogen.com/

COMMENT	VNTDATE|-13512298|

COMMENT	VNTDBDATE|-13512298|

COMMENT	LSOWNER|

COMMENT	VNTNAME|LHZ1014|

COMMENT	VNTAUTHORNAME|Demo User|

COMMENT	VNTOAUTHORNAME|Yao Yu|

FEATURES	Location/Qualifiers

CDS	complement(10627..11487)

/vntifkey="4"

/label=AmpR

rep_origin	9868..10456

/vntifkey="33"

/label=Rep(pMB1)

CDS	1893..3138

/vntifkey="4"

/label=ARS1

promoter	3139..3517

/vntifkey="29"

/label=TEF\Promoter

CDS	3518..4552

/vntifkey="4"

/label=HphMX4

terminator	4553..4787

/vntifkey="43"

/label=TEF\terminator

misc_structure 6208..6895

/standard_name="Tetrahymena telomere"

/citation=[5]

/vntifkey="88"

/label=Tetrahymena\telomere

misc_feature	complement(8723..9410)

/standard_name="Tetrahymena telomere"

/citation=[5]

/vntifkey="21"

/label=Tetrahymena\telomere

CDS	5348..6013

/vntifkey="4"

/label=TRP1\CDS

promoter	4788..5347

/vntifkey="29"

/label=TRP1\promoter

terminator	6014..6207

/vntifkey="43"

/label=TRP1\Terminator

CDS	1021..1686

/vntifkey="4"

/label=HIS3\CDS

promoter	419..1020

/vntifkey="29"

/label=HIS3\Promoter

terminator	1687..1886

/vntifkey="43"

/label=HIS3\Terminator

CDS	complement(6912..8714)

/vntifkey="4"

/label=Apu-GLA

BASE COUNT	3268 a	2536 c	2745 g	3138 t

ORIGIN

1 tcgcgcgttt cggtgatgac ggtgaaaacc tctgacacat gcagctcccg gagacggtca

61 cagcttgtct gtaagcggat gccgggagca gacaagcccg tcagggcgcg tcagcgggtg

121 ttggcgggtg tcggggctgg cttaactatg cggcatcaga gcagattgta ctgagagtgc

181 accatatgcg gtgtgaaata ccgcacagat gcgtaaggag aaaataccgc atcaggcgcc

241 attcgccatt caggctgcgc aactgttggg aagggcgatc ggtgcgggcc tcttcgctat

301 tacgccagct ggcgaaaggg ggatgtgctg caaggcgatt aagttgggta acgccagggt

361 tttcccagtc acgacgttgt aaaacgacgg ccagtgaatt gacgcgtatt gggatatcag

421 aatttttgtc tcttaccaaa aggatctacg tcaccattta aagccctttc agtcttctgc

481 ttcgcgtgca catgagaata tttcttctta aaccactcgt ccttatccat gccatctttt

541 tcccatgctt tcgttctctt ttgtatcggg aagttccttc caaatgttac tttcttgtct

601 ttgatgaagg attcatttgg gattttcacc gctggacgaa attctggtgg caaagcctct

661 ataaacagct ttcatgatat ttattgttga caacttaata ttactgttaa gaaaaggcgg

721 atgagaatac ctagctctct agtatctttc taatagatat accaagctca cattatctca

781 actcatctca tctcatctct atactgcaac tttttcttta gaaatttttc aagaagtgat

841 attttcttag acattttttt ttttttttaa aaggctccaa gaatgattca taatatgaaa

901 tcaagtataa atgcagctgt acaatggttt aacttagttt aagatccgtt cggaagtact

961 attattgtaa cattttacac aataagaagg tcttcaaata aggactttga taccgagaca

1021 atgacatacc cagaaaggaa ggcttttgtg tctagaataa caaatgagac aaaaattcag

1081 atagccatat ccttacatgg ggggcatatc tcaatcccaa attctatact ggatagaccg

1141 gagtcggacg ttgcaaaaca agctactggt tcacagatta ttgacattca aactggtatt

1201 ggatttctag atcatatgat tcatgcccta gcgaagcact ctggttggtc cttaattgtc

1261 gaatgtatcg gagatttgca tatcgatgac caccatacta cggaggattg tggtattgct

1321 ctaggacagg cttttaaaga agcattaggt catgtccgtg gtgtgagaag atttggtact

1381 ggatttgcac cattggacga agcattatca agggccgtcg ttgatctatc caacagacca

1441 ttcgccgtaa tagatttggg tttaaaaaga gaaaaaatcg gtgatctttc atgtgaaatg

1501 ataccacatt tcttggagtc atttgcggaa gctgcaagag taactttaca tgttgactgt

1561 ttaagaggct ttaatgatca tcacagaagt gagtcagcct ttaaggctct tgctgtggct

1621 atcagagagg ctatttccag taacggtaca aatgacgttc catctaccaa aggagttttg

1681 atgtaactgg ctactccata caaggcggta taaataaaat ataataacat gtttgtacaa

1741 taatgttttc ctatttatta cttcatatat tatatatgtt gcacctaaaa taaggaagtt

1801 attctcaaag ttattgtgta cttttatatt atttacatgg gaatttatat atatatatat

1861 gtgcgtgtgg tgacagcata acgcccctcg agatcgattg aagttttgtc caactatcca

1921 ctatggatat gcgttttgtt gattaacctt aaataacacg tatttcgcat tttccaaaag

1981 ccttttttca taactacaaa ctaactattt ttgtttattt tacatcagta aattatgcgc

2041 agaaaatatg taaggctata tactcaatat agtggaagag cctctggcta cttaatctgg

2101 gttcatatat tctgtcagtg gtatagtaaa tttaaatagt gaatttgggc gcatgtagag

2161 agatcttttg attaataacc cagtatttga ttcttgaatg ttatttgtct cttattaagt

2221 atttcagttt gatttatata ttttgaaagt aattgttgct ttaccttcca aacaataaaa

2281 aaatataaaa aaatgtaaaa aatataataa atattaaata aaatactact tgtttgaaaa

2341 tcagagaaaa tccaccaaaa tatcaatcat ttcaaggatt tccgaaccaa gttcgagata

2401 gtcctttaag gtcagtaaaa ttcagttgca cgtataacag atatttttca ttttgttcca

2461 atttaaaagt cccccatttt taaaattatt gaaaataaaa attaaaaaat taaaaggaat

2521 ctctctatgt cactttaaaa taaataaatt gaaaatgata tatcgttaaa agtcaaccgc

2581 aacaccacct atttctaaga ggagagttct aaaaaaatca tagtaccaca caggtaaact

2641 aaaacagcta aattcaacat aatacctgtg gatataatta catacaaaaa tataattaaa

2701 aaaatacatt aaaaataatt tatttttgta aaacccataa aatatatttt actttcggaa

2761 caacttttta acttataatt ttgttttaaa taaaaacgtt gtatttaaaa ataataaaat

2821 attaagtaaa aatttaaagc atatttattt aaaatataaa atactactaa aattactcta

2881 aacttcaaaa taaaaaaata ataaattata tagttttaaa attaataatt tgtatacacg

2941 tgaccagacc ataatagttt tcttttcttg aaactgccat gaatttaata gatttttttt

3001 acacatattt acagtaagtt ttgtttttac gttgaattaa tattgtatac gtacttagaa

3061 ggataacttc caactatgaa tatgtaggtt aaaaggtaaa tagagaagcc ttacttttat

3121 cagaaatgaa aggagctcag cttgccttgt ccccgccggg tcacccggcc agcgacatgg

3181 aggcccagaa taccctcctt gacagtcttg acgtgcgcag ctcaggggca tgatgtgact

3241 gtcgcccgta catttagccc atacatcccc atgtataatc atttgcatcc atacattttg

3301 atggccgcac ggcgcgaagc aaaaattacg gctcctcgct gcagacctgc gagcagggaa

3361 acgctcccct cacagacgcg ttgaattgtc cccacgccgc gcccctgtag agaaatataa

3421 aaggttagga tttgccactg aggttcttct ttcatatact tccttttaaa atcttgctag

3481 gatacagttc tcacatcaca tccgaacata aacaaccatg ggtaaaaagc ctgaactcac

3541 cgcgacgtct gtcgagaagt ttctgatcga aaagttcgac agcgtctccg acctgatgca

3601 gctctcggag ggcgaagaat ctcgtgcttt cagcttcgat gtaggagggc gtggatatgt

3661 cctgcgggta aatagctgcg ccgatggttt ctacaaagat cgttatgttt atcggcactt

3721 tgcatcggcc gcgctcccga ttccggaagt gcttgacatt ggggaattca gcggccgcga

3781 gagcctgacc tattgcatct cccgccgtgc acagggtgtc acgttgcaag acctgcctga

3841 aaccgaactg cccgctgttc tgcagccggt cgcggaggcc atggatgcga tcgctgcggc

3901 cgatcttagc cagacgagcg ggttcggccc attcggaccg caaggaatcg gtcaatacac

3961 tacatggcgt gatttcatat gcgcgattgc tgatccccat gtgtatcact ggcaaactgt

4021 gatggacgac accgtcagtg cgtccgtcgc gcaggctctc gatgagctga tgctttgggc

4081 cgaggactgc cccgaagtcc ggcacctcgt gcacgcggat ttcggctcca acaatgtcct

4141 gacggacaat ggccgcataa cagcggtcat tgactggagc gaggcgatgt tcggggattc

4201 ccaatacgag gtcgccaaca tcttcttctg gaggccgtgg ttggcttgta tggagcagca

4261 gacgcgctac ttcgagcgga ggcatccgga gcttgcagga tcgccgcggc tccgggcgta

4321 tatgctccgc attggtcttg accaactcta tcagagcttg gttgacggca atttcgatga

4381 tgcagcttgg gcgcagggtc gatgcgacgc aatcgtccga tccggagccg ggactgtcgg

4441 gcgtacacaa atcgcccgca gaagcgcggc cgtctggacc gatggctgtg tagaagtact

4501 cgccgatagt ggaaaccgac gccccagcac tcgtccgagg gcaaaggaat aatcagtact

4561 gacaataaaa agattcttgt tttcaagaac ttgtcatttg tatagttttt ttatattgta

4621 gttgttctat tttaatcaaa tgttagcgtg atttatattt tttttcgcct cgacatcatc

4681 tgcccagatg cgaagttaag tgcgcagaaa gtaatatcat gcgtcaatcg tatgtgaatg

4741 ctggtcgcta tactgctgtc gattcgatac taacgccgcc atccagtggt accgtcgcct

4801 tgagagaaat tagaagattc caaaaatcca ccgaactatt gatcagaaag ttgcctttcc

4861 aaagattggt tagagaaatc gcccaagact tcaagaccga tttgagattc caatcttctg

4921 ctatcggtgc cttgcaagaa tccgtcgaag cctacttggt ctccttgttc gaagacacca

4981 acttggctgc catccacgcc aagagagtca ccatccaaaa gaaggacatc aagttcgaag

5041 acaccaactt ggctgccatc cacgccaaga gagtcaccat ccaaaagaag gacatcaagt

5101 tggccagaag attgagaggt gagagatcgt gaatgttctt ttccccttcc tttccccttc

5161 ccttctaatt ttatctttat tatctactta ttttactggc tagtccagtt tttttcaacg

5221 cttcttttcc cctaggaaaa aatagagaag cgcaataagt atatcccagg tgtataatag

5281 tttaatatca atcgagtaat atacggtttt ctcaggaata acaactgcct tagtctagtc

5341 cacagacatg ctcgtcaaga tctgcggctt gcagtctgtt gaagctgctc aaacagcgct

5401 ggatcgcggc gcagacctgc tgggagtcat atgtgtcccc aacaggaaac gcaccgtcac

5461 gccagcaaca gcaaaacaaa tctcacaact ggttcaccag ggcaatcatt cccaggggaa

5521 tcaccaggcc aggctggtcg gggtgttccg gaaccagcct ctagaagaag tgctcgccct

5581 gtaccacgaa tacaacctag acgttatcca gcttcacggc aacgaagatg tggtccaatg

5641 gagaaaatgg attcccaagg acatcacgtt gatcaaggcg ttccagttcc ctggcgactg

5701 cgacgtggtg ttatcgccag cggtcgccca gctccagctg gaaaacgtgc tggtgctgtt

5761 cgattcgggc gaaggtggca cgggccagca gctcgactgg aacggcatgg ccagctggtg

5821 tcagaaccaa ggtactacca cccgcttcat actcgcggga ggactcaccc cagataacgt

5881 gggccacgcc atcacaagcc tcgcgcccca tgccatcgga gttgacgtca gcggaggtgt

5941 cgagacaaac ggccagaagg acatggccaa gatcgccgcc ttcatatcac aggcgagggg

6001 tctatctatc taataagtaa gtagagcatt ggttaatgaa tacataggta ataattactg

6061 tatttcctct agctagtacc ccgcagctca agcagaaccg gagatgaaga accacttgtc

6121 aatggacttg tcgattggct cgtctggcaa gacgtgtgct ggtggaacag cagcagcagc

6181 ttcggcagag taagtggcag tgtcgaccat ttttagataa aatttattaa tcatcattaa

6241 tttcttgaaa aacattttat ttattgatct tttataacaa aaaacccttc taaaagttta

6301 tttttgaatg aaaaacttat aaaaatttat gaaaactaca aaaaataaaa tttttaatta

6361 aaataatttt gataagaact tcaatctttg actagctagc ttagtcattt ttgagattta

6421 attaatattt tatgtttatt catatataaa ctattcaaaa tattatagaa tttaaacatt

6481 ttaacatctt aatcattcat aaataactaa aaatcaaagt attacatcaa taaataactt

6541 ttactcaatg tcaaagaatt attggggttg gggttggggt tggggttggg gttggggttg

6601 gggttggggt tggggttggg gttggggttg gggttggggt tggggttggg gttggggttg

6661 gggttggggt tggggttggg gttggggttg gggttggggt tggggttggg gttggggttg

6721 gggttggggt tggggttggg gttggggttg gggttggggt tggggttggg gttggggttg

6781 gggttggggt tggggttggg gttggggttg gggttggggt tggggttggg gtgggaaaac

6841 agcattcagg tattagaaga atatcctgat tcaggtgaaa atattgttga tgcgcgcgga

6901 tccgcaagct ttctccaaga gtcgttttca gtagcggtgc cggcacaatt accggtaacg

6961 gtgtaagaac ggtttgggtc agactcccaa gtgacggagc cgtcttgggc cttcttaatg

7021 tatttataat tgaaggaagt gccggtagcg aagttgatgg taacggtcca caatgggttg

7081 gaggaagtgt acttggaagc ggataaagcg acggcgttgg cggtattcca gttgcctaaa

7141 gctgggatgg agccgacgat gtagatgttc tcaccatagg aggtagtctt ttgttcgttg

7201 aaggtgacgg cgatagaggt tggagtagtg caagaaccgc cagtggtggt ggtaggagtg

7261 ccggtagatg gggaacccgg gttgccccag ttggtgttgg tggcggcagc acaaggacca

7321 gtagcagagc cagaagaaca agagttaggc aacttggcag aggaggcacc ccaagaagct

7381 ggcaaaacgt tggctctagc gttgtaagcg gtcaagaagg cagcgtagga ccaagttaag

7441 tcgacggcgg acaatggagt gccgtcggct ctgttgtact gctcagacaa ggcaccgttg

7501 gatggggtgt acttttgggc gatggacatg taggagtcgg cgtaggtttg aacggcgtta

7561 acgatggagg tgaaggtgac agtggaggaa gagtaagtgc cgacagcggc ggaggagtaa

7621 acatccttga agaaaggcaa ggagatggag gtgatggaga tggaaccgat cttcttccat

7681 tggtagacgg cgtcatataa ctgctcagcg gcggcgaagg tattcaagta ccatgggtta

7741 ccgttgtaat aagaatcttc tgggtatctg ccgacagcaa caccggaacc ttgagcaata

7801 ccttggttga tggagtagat ggaacggaaa gagtcagtga cgaccttgtg gttagccaaa

7861 gccttatcgg agcaaggctg gaaggtagtg gaatcacaag aggcggctgg atcgaagatg

7921 tggatggagg tcaaaatgga gttagcgtcc ttaccggatc taccaccgcc ggtattggac

7981 aaggcgtagg agccggtcca gtaggactgt aaaaagcaca agaccaatgg agcttgggaa

8041 acgcagtttg gacaagactt gcccaattgg gtggccaagt tgttaccctc gaccaaagca

8101 cggtactgaa cggcagtggt aaagaaggag gaggagttga tctcctccca taagtcaaag

8161 gtggtctggt tccagtactg agtgacgtag gacaagtcgt tttggacgat tggccagata

8221 atgttattaa cggtagtagt gttgccgtta gcgatcaagt aacgggagta agcgatcatg

8281 gcggtagctc tcaaagctgg accgtcacgt tgtggtctac cccaagcgcc agtgaaagga

8341 gtcaagtcaa cctcgaactt tggctcggct aagccaccgg aacacaaacc gccagatggg

8401 ttgttgacgg tttgtaactt agcttgagca gagatgtact gttggatcaa tggttccaag

8461 gacttgttgc cggcaatcaa ctgatcgacc aaggccttga agactaaagc ggagtcacgg

8521 gtccaagtgt agaagtagtc tgggttggac ttagatgggg aggcaacaac aacaccagcg

8581 gaagcgccgg aagccttaga accggaggcg ccgatgttgt ttaagacacc ttgcaaggcg

8641 acggtgtttt cggaagacaa ccaagaggat aaggagccgg tggctctttc ttgaatggac

8701 tctggagaag gtaaggatcc gcgcgcatca acaatatttt cacctgaatc aggatattct

8761 tctaatacct gaatgctgtt ttcccacccc aaccccaacc ccaaccccaa ccccaacccc

8821 aaccccaacc ccaaccccaa ccccaacccc aaccccaacc ccaaccccaa ccccaacccc

8881 aaccccaacc ccaaccccaa ccccaacccc aaccccaacc ccaaccccaa ccccaacccc

8941 aaccccaacc ccaaccccaa ccccaacccc aaccccaacc ccaaccccaa ccccaacccc

9001 aaccccaacc ccaaccccaa ccccaacccc aaccccaacc ccaaccccaa ccccaataat

9061 tctttgacat tgagtaaaag ttatttattg atgtaatact ttgattttta gttatttatg

9121 aatgattaag atgttaaaat gtttaaattc tataatattt tgaatagttt atatatgaat

9181 aaacataaaa tattaattaa atctcaaaaa tgactaagct agctagtcaa agattgaagt

9241 tcttatcaaa attattttaa ttaaaaattt tattttttgt agttttcata aatttttata

9301 agtttttcat tcaaaaataa acttttagaa gggttttttg ttataaaaga tcaataaata

9361 aaatgttttt caagaaatta atgatgatta ataaatttta tctaaaaatg gtcgacctgc

9421 aggcatgcaa gcttccaatg gcgcgccgag cttggcgtaa tcatggtcat agctgtttcc

9481 tgtgtgaaat tgttatccgc tcacaattcc acacaacata cgagccggaa gcataaagtg

9541 taaagcctgg ggtgcctaat gagtgagcta actcacatta attgcgttgc gctcactgcc

9601 cgctttccag tcgggaaacc tgtcgtgcca gctgcattaa tgaatcggcc aacgcgcggg

9661 gagaggcggt ttgcgtattg ggcgctcttc cgcttcctcg ctcactgact cgctgcgctc

9721 ggtcgttcgg ctgcggcgag cggtatcagc tcactcaaag gcggtaatac ggttatccac

9781 agaatcaggg gataacgcag gaaagaacat gtgagcaaaa ggccagcaaa aggccaggaa

9841 ccgtaaaaag gccgcgttgc tggcgttttt ccataggctc cgcccccctg acgagcatca

9901 caaaaatcga cgctcaagtc agaggtggcg aaacccgaca ggactataaa gataccaggc

9961 gtttccccct ggaagctccc tcgtgcgctc tcctgttccg accctgccgc ttaccggata

10021 cctgtccgcc tttctccctt cgggaagcgt ggcgctttct catagctcac gctgtaggta

10081 tctcagttcg gtgtaggtcg ttcgctccaa gctgggctgt gtgcacgaac cccccgttca

10141 gcccgaccgc tgcgccttat ccggtaacta tcgtcttgag tccaacccgg taagacacga

10201 cttatcgcca ctggcagcag ccactggtaa caggattagc agagcgaggt atgtaggcgg

10261 tgctacagag ttcttgaagt ggtggcctaa ctacggctac actagaagaa cagtatttgg

10321 tatctgcgct ctgctgaagc cagttacctt cggaaaaaga gttggtagct cttgatccgg

10381 caaacaaacc accgctggta gcggtggttt ttttgtttgc aagcagcaga ttacgcgcag

10441 aaaaaaagga tctcaagaag atcctttgat cttttctacg gggtctgacg ctcagtggaa

10501 cgaaaactca cgttaaggga ttttggtcat gagattatca aaaaggatct tcacctagat

10561 ccttttaaat taaaaatgaa gttttaaatc aatctaaagt atatatgagt aaacttggtc

10621 tgacagttac caatgcttaa tcagtgaggc acctatctca gcgatctgtc tatttcgttc

10681 atccatagtt gcctgactcc ccgtcgtgta gataactacg atacgggagg gcttaccatc

10741 tggccccagt gctgcaatga taccgcgaga cccacgctca ccggctccag atttatcagc

10801 aataaaccag ccagccggaa gggccgagcg cagaagtggt cctgcaactt tatccgcctc

10861 catccagtct attaattgtt gccgggaagc tagagtaagt agttcgccag ttaatagttt

10921 gcgcaacgtt gttgccattg ctacaggcat cgtggtgtca cgctcgtcgt ttggtatggc

10981 ttcattcagc tccggttccc aacgatcaag gcgagttaca tgatccccca tgttgtgcaa

11041 aaaagcggtt agctccttcg gtcctccgat cgttgtcaga agtaagttgg ccgcagtgtt

11101 atcactcatg gttatggcag cactgcataa ttctcttact gtcatgccat ccgtaagatg

11161 cttttctgtg actggtgagt actcaaccaa gtcattctga gaatagtgta tgcggcgacc

11221 gagttgctct tgcccggcgt caatacggga taataccgcg ccacatagca gaactttaaa

11281 agtgctcatc attggaaaac gttcttcggg gcgaaaactc tcaaggatct taccgctgtt

11341 gagatccagt tcgatgtaac ccactcgtgc acccaactga tcttcagcat cttttacttt

11401 caccagcgtt tctgggtgag caaaaacagg aaggcaaaat gccgcaaaaa agggaataag

11461 ggcgacacgg aaatgttgaa tactcatact cttccttttt caatattatt gaagcattta

11521 tcagggttat tgtctcatga gcggatacat atttgaatgt atttagaaaa ataaacaaat

11581 aggggttccg cgcacatttc cccgaaaagt gccacctgac gtctaagaaa ccattattat

11641 catgacatta acctataaaa ataggcgtat cacgaggccc tttcgtc

//
