## Supplementary Table S3 for "Transfer of disulfide bond formation modules via yeast artificial chromosomes promotes the expression of heterologous proteins in *Kluyveromyces marxianus*"

LOCUS	LHZ1015	10276 bp	DNA	circular	23-FEB-2022

SOURCE

ORGANISM

COMMENT	This file is created by Vector NTI

http://www.invitrogen.com/

COMMENT	VNTDATE|-14275849|

COMMENT	VNTDBDATE|-14277271|

COMMENT	LSOWNER|

COMMENT	VNTNAME|LHZ1015|

COMMENT	VNTAUTHORNAME|Demo User|

COMMENT	VNTOAUTHORNAME|Yao Yu|

FEATURES	Location/Qualifiers

CDS	complement(9216..10076)

/vntifkey="4"

/label=AmpR

rep_origin	8457..9045

/vntifkey="33"

/label=Rep(pMB1)

CDS	1893..3138

/vntifkey="4"

/label=ARS1

promoter	3139..3517

/vntifkey="29"

/label=TEF\Promoter

CDS	3518..4552

/vntifkey="4"

/label=HphMX4

terminator	4553..4787

/vntifkey="43"

/label=TEF\terminator

CDS	5348..6013

/vntifkey="4"

/label=TRP1\CDS

promoter	4788..5347

/vntifkey="29"

/label=TRP1\promoter

terminator	6014..6207

/vntifkey="43"

/label=TRP1\Terminator

CDS	1021..1686

/vntifkey="4"

/label=HIS3\CDS

promoter	419..1020

/vntifkey="29"

/label=HIS3\Promoter

terminator	1687..1886

/vntifkey="43"

/label=HIS3\Terminator

misc_feature	6208..6841

/vntifkey="21"

/label=KM\Telomere

CDS	6858..7357

/vntifkey="4"

/label=5'Abe-GLA

misc_feature	complement(7366..7999)

/vntifkey="21"

/label=KM\Telomere

BASE COUNT	2881 a	2294 c	2243 g	2858 t

ORIGIN

1 tcgcgcgttt cggtgatgac ggtgaaaacc tctgacacat gcagctcccg gagacggtca

6181 ttcggcagag taagtggcag tgtcgacggt gtacggattt gattagttat gtggtgtacg

6241 gatttgatta gttatgtggt gtacggattt gattagttat gtggtgtacg gatttgatta

6301 gttatgtggt gtacggattt gattagttat gtggtgtacg gatttgatta gttatgtggt

6361 gtacggattt gattagttat gtggtgtacg gatttgatta gttatgtggt gtacggattt

6421 gattagttat gtggtgtacg gatttgatta gttatgtggt gtacggattt gattagttat

6481 gtggtgtacg gatttgatta gttatgtggt gtacggattt gattagttat gtggtgtacg

6541 gatttgatta gttatgtggt gtacggattt gattagttat gtggtgtacg gatttgatta

6601 gttatgtggt gtacggattt gattagttat gtggtgtacg gatttgatta gttatgtggt

6661 gtacggattt gattagttat gtggtgtacg gatttgatta gttatgtggt gtacggattt

6721 gattagttat gtggtgtacg gatttgatta gttatgtggt gtacggattt gattagttat

6781 gtggtgtacg gatttgatta gttatgtggt gtacggattt gattagttat gtggtgtacg

6841 ggcggatccg caagcttttc caagttagat tcaagccttc cgaggacacc gccttagaca

6901 ctgtcgatga cggcaccttg cagtccttgt tggacaacat cggcttgaac ggttctaatg

6961 cttgggacac cagaccgggt ttggttatcg cctccccttc caagaaggac cctaactact

7021 tcttcacttg gactcgtgac tccgccttgg tcttaaaatg tatcaccgac gctttcgctg

7081 ctggtaatac cgccttgcaa gaaaccatcc acgagtacat ctcttcccaa gctcgtatcc

7141 agttattgaa caccagatcc ggcggtttgt cctccggcgg tttaggcgag ccaaagtacc

7201 gtgttgacga gaccccttac aacgaagatt ggggtagacc acaagctgat ggtccagcct

7261 tgcgtgccac tgccttgatt gcttacgccc gttggttatt ggaaaatgac tactacgacg

7321 ttgccaagtc tatcgtttgg ccagttgtta agaacgagga tccgcccgta caccacataa

7381 ctaatcaaat ccgtacacca cataactaat caaatccgta caccacataa ctaatcaaat

7441 ccgtacacca cataactaat caaatccgta caccacataa ctaatcaaat ccgtacacca

7501 cataactaat caaatccgta caccacataa ctaatcaaat ccgtacacca cataactaat

7561 caaatccgta caccacataa ctaatcaaat ccgtacacca cataactaat caaatccgta

7621 caccacataa ctaatcaaat ccgtacacca cataactaat caaatccgta caccacataa

7681 ctaatcaaat ccgtacacca cataactaat caaatccgta caccacataa ctaatcaaat

7741 ccgtacacca cataactaat caaatccgta caccacataa ctaatcaaat ccgtacacca

7801 cataactaat caaatccgta caccacataa ctaatcaaat ccgtacacca cataactaat

7861 caaatccgta caccacataa ctaatcaaat ccgtacacca cataactaat caaatccgta

7921 caccacataa ctaatcaaat ccgtacacca cataactaat caaatccgta caccacataa

7981 ctaatcaaat ccgtacaccg tcgacctgca ggcatgcaag cttccaatgg cgcgccgagc

8041 ttggcgtaat catggtcata gctgtttcct gtgtgaaatt gttatccgct cacaattcca

8101 cacaacatac gagccggaag cataaagtgt aaagcctggg gtgcctaatg agtgagctaa

8161 ctcacattaa ttgcgttgcg ctcactgccc gctttccagt cgggaaacct gtcgtgccag

8221 ctgcattaat gaatcggcca acgcgcgggg agaggcggtt tgcgtattgg gcgctcttcc

8281 gcttcctcgc tcactgactc gctgcgctcg gtcgttcggc tgcggcgagc ggtatcagct

8341 cactcaaagg cggtaatacg gttatccaca gaatcagggg ataacgcagg aaagaacatg

8401 tgagcaaaag gccagcaaaa ggccaggaac cgtaaaaagg ccgcgttgct ggcgtttttc

8461 cataggctcc gcccccctga cgagcatcac aaaaatcgac gctcaagtca gaggtggcga

8521 aacccgacag gactataaag ataccaggcg tttccccctg gaagctccct cgtgcgctct

8581 cctgttccga ccctgccgct taccggatac ctgtccgcct ttctcccttc gggaagcgtg

8641 gcgctttctc atagctcacg ctgtaggtat ctcagttcgg tgtaggtcgt tcgctccaag

8701 ctgggctgtg tgcacgaacc ccccgttcag cccgaccgct gcgccttatc cggtaactat

8761 cgtcttgagt ccaacccggt aagacacgac ttatcgccac tggcagcagc cactggtaac

8821 aggattagca gagcgaggta tgtaggcggt gctacagagt tcttgaagtg gtggcctaac

8881 tacggctaca ctagaagaac agtatttggt atctgcgctc tgctgaagcc agttaccttc

8941 ggaaaaagag ttggtagctc ttgatccggc aaacaaacca ccgctggtag cggtggtttt

9001 tttgtttgca agcagcagat tacgcgcaga aaaaaaggat ctcaagaaga tcctttgatc

9061 ttttctacgg ggtctgacgc tcagtggaac gaaaactcac gttaagggat tttggtcatg

9121 agattatcaa aaaggatctt cacctagatc cttttaaatt aaaaatgaag ttttaaatca

9181 atctaaagta tatatgagta aacttggtct gacagttacc aatgcttaat cagtgaggca

9241 cctatctcag cgatctgtct atttcgttca tccatagttg cctgactccc cgtcgtgtag

9301 ataactacga tacgggaggg cttaccatct ggccccagtg ctgcaatgat accgcgagac

9361 ccacgctcac cggctccaga tttatcagca ataaaccagc cagccggaag ggccgagcgc

9421 agaagtggtc ctgcaacttt atccgcctcc atccagtcta ttaattgttg ccgggaagct

9481 agagtaagta gttcgccagt taatagtttg cgcaacgttg ttgccattgc tacaggcatc

9541 gtggtgtcac gctcgtcgtt tggtatggct tcattcagct ccggttccca acgatcaagg

9601 cgagttacat gatcccccat gttgtgcaaa aaagcggtta gctccttcgg tcctccgatc

9661 gttgtcagaa gtaagttggc cgcagtgtta tcactcatgg ttatggcagc actgcataat

9721 tctcttactg tcatgccatc cgtaagatgc ttttctgtga ctggtgagta ctcaaccaag

9781 tcattctgag aatagtgtat gcggcgaccg agttgctctt gcccggcgtc aatacgggat

9841 aataccgcgc cacatagcag aactttaaaa gtgctcatca ttggaaaacg ttcttcgggg

9901 cgaaaactct caaggatctt accgctgttg agatccagtt cgatgtaacc cactcgtgca

9961 cccaactgat cttcagcatc ttttactttc accagcgttt ctgggtgagc aaaaacagga

10021 aggcaaaatg ccgcaaaaaa gggaataagg gcgacacgga aatgttgaat actcatactc

10081 ttcctttttc aatattattg aagcatttat cagggttatt gtctcatgag cggatacata

10141 tttgaatgta tttagaaaaa taaacaaata ggggttccgc gcacatttcc ccgaaaagtg

10201 ccacctgacg tctaagaaac cattattatc atgacattaa cctataaaaa taggcgtatc

10261 acgaggccct ttcgtc

//
